# Clearing truncated tau protein restores neuronal function and prevents microglia activation in tauopathy mice

**DOI:** 10.1101/2024.05.21.595198

**Authors:** Alejandro Martín-Ávila, Swananda R. Modak, Hameetha B. Rajamohamedsait, Andie Dodge, Dov B. Shamir, Senthilkumar Krishnaswamy, Leslie A. Sandusky-Beltran, Marilyn Walker, Yan Lin, Erin E. Congdon, Einar M. Sigurdsson

## Abstract

Tau protein truncated at aspartate 421 (Asp421) is a characteristic feature of Alzheimer’s disease (AD) and other tauopathies. It is likely to have a role in their pathogenesis by promoting tau aggregation. Here, using two tauopathy mouse models, we show that a monoclonal antibody against Asp421, 5G2, led to a) a 59-74% clearance of insoluble tau protein in the brains of JNPL3 tauopathy mice following a thirteen-week treatment period, b) a 46% decrease of tau levels in brain interstitial fluid immediately following a single dose of 5G2 as examined by brain microdialysis in awake JNPL3 mice, c) improved neuronal function and d) reduced microglial activation as determined by two-photon imaging in awake PS19 tauopathy mice, where we also found tau accumulation earlier than signs of microglial activation. For mechanistic insight using culture models, 5G2 prevented toxicity of AD brain-derived pathological tau protein, cleared intracellular tau, and prevented microgliosis. We also knocked down the intracellular Fc receptor and ubiquitin E3 ligase, TRIM21, and found a reduction in cellular retention of tau antibodies, which appeared to reduce the acute efficacy (24 h) of tau antibodies but not their longer-term efficacy (5 days). Overall, these findings strongly support the feasibility of targeting Asp421 truncated tau protein to treat tauopathies, indicate that tau-associated abnormalities of neuronal activity precede microglial activation and that antibody-mediated tau clearance via the TRIM21 pathway is mostly transient.

## INTRODUCTION

Tauopathies are defined by the hyperphosphorylation, truncation, sometimes mutation, and aggregation of the tau protein, which is a cytoskeletal protein that is primarily found in neurons. Alzheimer’s disease (AD) is the most common one by far but is also characterized by amyloid-β deposition. Of the primary tauopathies, progressive supranuclear palsy (PSP) has the largest number of subjects. In recent years, immunotherapies targeting the tau protein have advanced from proof-of-concept animal studies to clinical trials (Asuni et al., 2006;Asuni et al., 2007;Boutajangout et al., 2010a;Boutajangout et al., 2011;Congdon et al., 2023;Sigurdsson, 2024). Currently, there are nine antibodies and two tau immunogens being examined in Phase 1-3 trials in patients with AD or in healthy subjects. The epitopes that are being targeted are different hyperphosphorylated forms of tau, unphosphorylated tau, and misfolded tau.

In addition to these epitopes, various truncations of the tau protein have been detected in AD, other tauopathies and related animal models. These truncated forms may promote tau aggregation and/or toxicity (Chun and Johnson, 2007;Cotman et al., 2005). Of these, tau truncated at aspartate 421 is the most prominent and is primarily thought to be the result of caspase-3 cleavage. It has been reported in different animal models to occur late or early in the progression of tau pathology (de Calignon A. et al., 2010;Delobel et al., 2008), and has recently been suggested to be an early epitope based on detailed proteomic analysis of AD brain tissue at different stages of the disease (Wesseling et al., 2020).

We have previously shown that acute anti-tau antibody treatment clears pathological tau, improves cognition, and targeting phosphorylated tau reverses functional abnormalities in L2/3 pyramidal neurons in the cortex of the JNLP3 mice (Congdon et al., 2016;Wu et al., 2018;Wu et al., 2020) but targeting truncated tau for therapeutic clearance has not been well examined. We have previously reported our preliminary findings showing the efficacy of targeting the most prominent tau truncation at aspartate 421 in different culture models (Modak et al., 2015;Modak and Sigurdsson, 2017).

Other investigators have seen similar effects in other culture models with another Asp421 targeting antibody (Nicholls et al., 2017;Nobuhara et al., 2017). Most of pathological tau protein resides within neurons but a small portion is found extracellularly, and it may be taken up into adjacent neurons, where it may seed further tau aggregation (Clavaguera et al., 2017;Gibbons et al., 2019;Vaquer-Alicea and Diamond, 2019). The majority of the preclinical studies examining the therapeutic potential of tau antibodies have not analyzed if the antibodies are working intra- or extracellularly. Of those that do, some are found intraneuronally (Asuni et al., 2007;Chandupatla et al., 2020;Collin et al., 2014;Congdon et al., 2013;Congdon et al., 2016;Congdon et al., 2019;Gu et al., 2013;Kondo et al., 2015;Krishnamurthy et al., 2011;Krishnaswamy et al., 2014;McEwan et al., 2017;Mukadam et al., 2023;Shamir et al., 2016;Shamir et al., 2020;Wu et al., 2018;Zhang et al., 2024), whereas others do not appear to be taken up (Castillo-Carranza et al., 2014;d’Abramo et al., 2015;Yanamandra et al., 2015). These differences may be explained by varying antibody charge, which we have shown to influence neuronal uptake of tau antibodies (Congdon et al., 2019).

With regard to the pathways involved in antibody-mediated intracellular clearance of tau, we and others have repeatedly shown the involvement of the endosomal-lysosomal system, in which various tau antibodies associate with pathological tau aggregates after uptake (for review see (Congdon et al., 2022;Sandusky-Beltran and Sigurdsson, 2020). Other investigators have shown the involvement of the proteasomal pathway via ubiquitin E3 ligase named TRIM21 (McEwan et al., 2017;Mukadam et al., 2023). To date, most of the therapeutic studies on tau antibodies have been performed in animals or in neuronal cultures. Possible involvement of glial cells, in particular microglia in this context is not well known, and conflicting findings have been reported (Andersson et al., 2019;Lee et al., 2016;Luo et al., 2015;Zilkova et al., 2020).

Microglia activation has been reported in Alzheimer’s disease (AD), tauopathies, and related models (Davidson et al., 2023;Mary et al., 2024). These cells may have beneficial and/or detrimental effects on neurons depending on the stage of neurodegeneration but much is unknown regarding their relationship with the progression of neurodegeneration.

To study these different variables, we treated with Asp421 antibodies primary tauopathy mouse neurons, primary mixed tauopathy mouse cortical cultures and differentiated human neuroblastoma cells, all of which had been fed pathological tau proteins derived from Alzheimer’s brains. We also knocked down TRIM21 in the differentiated human culture to study the potential involvement of that pathway in antibody-mediated tau clearance and blocking of tau toxicity. One of the antibodies was more effective in the different culture models, presumably because of its higher affinity for the truncated Asp421 tau. Their individual efficacies were similar in models with or without microglia, indicating limited if any involvement of microglia. However, antibody efficacy was associated with a decrease in a microglial marker, suggesting an anti-inflammatory effect that is likely to be related to tau clearance. Notably, TRIM21 knockdown diminished antibody retention within the neurons but primarily reduced their efficacy under acute (24 h) but not long-term (5 days) conditions. Finally, in vivo efficacy of the more effective antibody was confirmed in three studies in two tauopathy mouse models that showed its acute tau clearance, improvement in neuronal function and reduced microglial activation as well as its clearance of insoluble tau following chronic treatment. Overall, these findings support the therapeutic potential of targeting Asp421 truncated tau protein, and provide important insight into the mechanisms involved.

## MATERIALS AND METHODS

### Antibody generation

Monoclonal antibodies were generated by GenScript Inc. (Paramus, NJ). Wild type (WT) BALB/c mice were immunized with a peptide corresponding to tau region 407-421 (cHLSNVSSTGSIDMVD). The peptide was conjugated to keyhole limpet hemocyanin via the cysteine residue, and mice showing a satisfactory immune response were used in hybridoma production. Monoclonal antibodies 5G2 and 1G11 were selected and endotoxin purified from the culture supernatant.

### Antibody characterization

#### Surface plasmon resonance

The binding kinetics of 5G2 and 1G11 to tau peptides encompassing amino acids 407-421 and 407-423 were undertaken through Biacore 2000 (GE Healthcare) using surface plasmon resonance analysis according to the manufacturer’s protocol. The 407-421 peptide consists of the immunogen, whereas the longer 407-423 peptide was used to determine if the antibodies bound to a different epitope than the truncated free Asp421 terminus. Both 5G2 and 1G11, 10 µg/ml each, were diluted in 10 mM sodium acetate, pH 5.0 and immobilized on a separate CM5 sensor chip with an amine coupling kit (7 min contact time at 5 µl/min flow rate). One channel of each sensor chip was prepared in the same way without antibodies and was used to detect non-specific binding of the peptide. The HBS-EP buffer containing 10 mM HEPES, pH 7.4, 150 mM NaCl, 3.4 mM EDTA, and 0.005% surfactant P20 was used at a flow rate of 5 µl/min at 25°C. This was followed by regenerating the chip surface with 10 µl of buffer containing 500 mM NaCl and 0.1 M glycine HCl, pH 8.0 after undertaking each measurement. Binding of 5G2 and 1G11 antibodies to the Asp421 epitope was determined by calculating the equilibrium dissociation constant (K_D_) using BIA evaluation software with K_D =_ K_off_/K_on_.

#### Immunohistochemistry

Brain tissue staining using the two tau antibodies, 5G2 and 1G11, targeting truncated tau was conducted on mouse and human brains, and compared to PHF1 staining. The mouse brains were from JNPL3 tauopathy mice and wild-type controls of the same strain background and the brain processing and staining was conducted on free floating fixed coronal brain sections (40 µm) as we have described previously in detail (Rajamohamedsait and Sigurdsson, 2012), with the primary antibodies incubated at 1:1000 dilution (5G2 and 1G11 1 mg/ml; PHF1 cell culture supernatant) overnight at 4°C. For the human brain tissue, paraffin embedded brain block from the frontal pole was sectioned (8 µm) and subsequently mounted on slides. All the sections were deparaffinized in 3 changes of xylene and descending ethanol series (100% and 95%) for 5 min each followed by washing in running distilled water for a further 5 min. Antigen retrieval was undertaken in 88% formic acid for 7 min followed by incubating the sections in boiling citrate buffer (10 mM, pH 6.0) for 17 min. The endogenous peroxidase activity was quenched by incubating the sections in 0.3% hydrogen peroxidase for 30 min and subsequently washed with TBS twice for 5 min. As for the mouse tissue, the human sections were incubated overnight at 4°C with the primary antibodies. This was followed by washing the sections twice in TBS for 5 min each followed by incubating in the Biocare Multimer HRP secondary antibody (Biocare Medical Pacheco CA) for 1 h at room temperature. This was followed with subsequent washes with TBS twice for 5 min each and then with 0.2 M sodium acetate for further 10 min. The chromogenic reaction was developed in DAB (3, 3’-diaminobenzidine tetrahydrochloride) and nickel ammonium sulfate in 0.003% H_2_O_2_ and 0.2 M sodium acetate, analogous to the mouse tissue staining. The sections turned brown, after which the chromogenic reaction was terminated by washing the sections in 0.2 M sodium acetate and then TBS for 10 min each. This was followed by air drying the sections for 24 h, dehydrating and defatting in series of ethanol (70-100%), followed by Citrisolv and then cover slipped using Depex mounting media (BDH Laboratory Supplies).

### PHF preparation

Paired helical filament (PHF) enriched tau protein was prepared from human AD brain slices similar to as previously described (Lee et al., 1999). The tissue was homogenized in a buffer containing 0.75 M NaCl, 1 mM EGTA, 0.5 mM MgSO_4_, and 100 mM 2-(N-morpholino) ethanesulfonic acid, pH 6.5, with protease inhibitor cocktail (Roche, Indianapolis, IN) and centrifuged at 11, 000 x g for 20 min at 4°C.The resulting low speed supernatant was then incubated with 1% sarkosyl for one hour at room temperature. This mixture was then centrifuged at 100, 000 x g for 60 minutes. The supernatant was removed, and the pellet washed with 1% sarkosyl. The tau was then resolubilized by being briefly heated to 37°C in 50 mM Tris-HCl buffer (pH 7.4) using 0.5 mL of buffer for each mg of initial weight of brain sample protein, and then dialyzed in PBS overnight using a 3500 MW cassette. This resulted in PHF-enriched fraction as verified by electron microscopy that was used in the culture studies to promote tau toxicity and seeding.

### Mixed cortical cultures and primary neurons

Mixed cortical cultures were prepared from postnatal Day 0 JNPL3 pups. The 24 well plates were coated with poly-L-Lysine for 3 h in the incubator with 5% CO_2_ at 37°C. After 3 h, the plates were washed with HBSS+++ buffer (975 ml Hank’s balanced salt solution, 10 ml of 1 M HEPES, 5 ml of penicillin/streptomycin (P/S) and 10 ml of 100 mM sodium pyruvate), followed by addition of plating media (443.5 ml DMEM, 50 ml FBS, 2.5 ml P/S and 4 ml B27). The plates were kept in the incubator until the brains were harvested after the removal of meninges and brainstem. The brain tissue was washed 5 times in HBSS+++ buffer and then incubated with 200 µl of 0.05% trypsin for 15 min. This was followed by addition of equal volume of plating media to neutralize the effect of trypsin. The tissue was washed 5 times with HBSS+++ buffer and centrifuged for 1 min at 0.5 x g at room temperature. The tissue was resuspended in 2 ml of plating media, mechanically dissociated and then plated. The cells were incubated for 5-6 days to develop the neuronal processes, which was followed by treating the cells with PHF and antibodies. For primary neurons, the plating media was replaced by neurobasal media (499 ml Neurobasal A, 1 ml B27 and 17 µl of basal medium eagle) the following day and maintained for the next 5 days to develop processes.

### Differentiation of SH-SY5Y cells

SH-SY5Y human neuroblastoma cells were obtained from ATCC. The cells were maintained in DMEM media with GlutaMAX (Invitrogen) supplemented with 10% heat inactivated FBS, 10, 000 Units/mL Penicillin and 10, 000 μg/mL streptomycin. The cells were plated at 4x10^2^ cells/mm^2^ and then grown for 3-5 days to reach about 70% confluency before starting their differentiation. The cells were double-differentiated using retinoic acid (RA) and BDNF (brain-derived neurotrophic factor). For differentiation, the cells were maintained in the media containing 10 µM RA (Sigma Aldrich) and 1% FBS for 5 days. This was followed by washing the cells twice in DMEM media and then treating with 50 ng/ml BDNF (Alomone Labs) in serum free media for the next 2 days. For all the experiments, SH-SY5Y cells were grown in the media supplemented with only BDNF. For the TRIM21 studies, both the naïve differentiated and TRIM21 KD (knock down) differentiated SH-SY5Y cells were pretreated with PHF (5 µg/ml) for 24 h followed by treatment with 5 µg/ml tau antibodies (5G2, 1G11 and 4E6) for further 24 h and 5 days. This was followed by undertaking Western blots against total tau and GAPDH.

### Transfection of TRIM21 shRNA plasmid into SH-SY5Y cells

Plasmid containing the TRIM21 shRNA target sequence GAGTTGGCTGAGAAGTTGGAA with a pLKO.1-hPGK-Puro-CMV-tGFP vector (Sigma, TRCN0000010839, Clone ID NM_003141.x-555s1c1) was transfected into SH-SY5Y cells using Lipofectamine 3000 (Invitrogen) according to the manufacturer’s instructions. Cells were transfected for 3 days, selected using up to 1 µg/mL puromycin in complete media, and further enriched using flow cytometry cell sorting (Moflo XDP, Beckman Coulter) for GFP positive cells.

### Brain extraction for protein preparation

At the end of each animal study, the animals were deeply anesthetized with ketamine/xylazine ((250 mg/50 mg per kg body weight, i.p.), perfused with PBS and the brain extracted.

### Western blots

Western blots were conducted for the samples from mixed cortical cultures, primary neurons, differentiated SH-SY5Y cells and mouse brains. The cell lysates or brain were homogenized in RIPA buffer containing 50 mM Tris-HCl, pH 7.4, 150 mM NaCl, 1 mM EDTA, 1 mM NaF, 1 mM Na_3_VO_4_, and 1 µg/ml cOmplete^TM^ protease inhibitor mixture (Roche). After incubation on ice for 15 min, the lysate was sonicated twice with 1 min rest on ice between the two pulses. This was followed by incubation on ice for additional 15 min and centrifugation at 13, 000 x g for 5 min at 4°C to remove membrane fraction. The supernatant was collected and total protein concentration was measured using BCA assay (Thermo Scientific). Samples were diluted in O+ buffer (62.5 mM Tris-HCl, pH 6.8, 5% glycerol, 2-mercaptoethanol, 2.3% SDS, 1 mM EGTA, 1 mM phenylmethylsulfonyl fluoride, 1 mM Na_3_VO_4_, and 1 µg/ml cOmplete protease inhibitor mixture), boiled for 5 min and loaded onto a 10% SDS-PAGE gel. Gels were transferred to nitrocellulose membranes at 100 V for 1 h and blocked in 5% dried milk in TBS-T. The immunoblots were then probed for total tau (1:1000, Dako rabbit), pSer199 (1:1000, Life Technologies, rabbit), NeuN (1:1000, Millipore, rabbit), GAPDH (1:5000, Abcam, rabbit), Iba1 (1:1000, Santa Cruz, rabbit) and TRIM21 (Ro/SSA) (1:1000, Santa Cruz, mouse) in Superblock ^TM^ blocking buffer in TBS (Thermo Fisher Scientific) overnight at 4°C. The blots were then washed and probed with anti-horseradish peroxidase (HRP) conjugated mouse or rabbit secondary antibody for 1 h followed by washing the blots 3 times with TBS-T. ECL substrate (Thermo Fisher Scientific) was used to detect the signal and the chemiluminescent signal was quantified using Fuji LAS-4000 image analyzer. For antibody uptake studies, the primary antibody incubation was bypassed with the addition of anti-mouse IgG1 HRP conjugated secondary antibody (Thermo Fisher Scientific), which was followed by further quantification of signal using Fuji LAS-4000 image analyzer.

### Animals

Cx3cr1^GPF^ (JAX # 005582), Cx3cr1^CreER^ (JAX # 021160), Thy-1^GCaMP6^, flox-tdTomato (JAX # 007909), PS19 (JAX # 008169) and JNLP3 (Lewis et al., 2000) mice were used in this study. Cx3cr1^GPF^, Cx3cr1^CreER^, and flox-tdTomato were crossed (heterozygous to heterozygous) with the PS19 tauopathy mouse model to visualize cortical microglia. Thy-1^GCaMP6^ mice were crossed (heterozygous to heterozygous) with the PS19 mice to visualize L2/3 pyramidal neurons. Thy-1^GCaMP6^ mice were engineered at New York University and maintained on a C57BL/6 background (Cichon et al., 2020). Mice were bred and housed in the animal facility at NYU Langone Medical Center, with 12h/12h light/dark cycle and ad libitum access to food and water.

Tamoxifen (Sigma) was given as a solution in corn oil to adult Cx3cr1^CreER^: tdTomato^flox^ mice by gavage to recombine tdTomato structural marker into microglia. Animals received two doses of 10 mg of tamoxifen 48 hours apart 30 days before experimentation.

### Chronic tau antibody treatment

For passive immunization study, 7-8 months old homozygous female JNPL3 mice were enrolled (Taconic). In total, the 22 JNPL3 mice that were enrolled in the study were segregated into treatment group that received 5G2 vaccine (n=11) and control group that was administered pooled IgG (Equitech-Bio Inc., Kerrville, TX, USA); n=11). The antibody was injected intraperitoneally at 10 mg/kg (250 µg/125 µl for 25 g weight). Thirteen weekly injections were administered. This is the same experimental design as in our original passive immunization study with the PHF1 antibody (Boutajangout et al., 2011) that was based on prior Aβ antibody studies (Schenk et al., 1999). Analysis at the end of the study revealed that one of the treated mouse did not express tau although it contained the transgene (see CP27 negative lanes in Western blots, Figure 3). It was therefore excluded from the study.

The brain tissue was homogenized in (5x vol/w) modified radioimmunoprecipitation assay (RIPA) buffer (50 mM Tris-HCl, 150 mM NaCl, 1 mM ethylene diamine tetra-acetic acid (EDTA), 1% Nonidet P-40, pH 7.4) containing protease and phosphatase inhibitors (4 nM phenylmethylsulfonyl fluoride (PMSF), 1 mM NaF, 1 mM Na_3_VO_4_, 1× cOmplete protease inhibitor cocktail (Roche, Indianapolis, IN, USA), and 0.25% sodium deoxycholate). The samples were then subjected to low-speed spin at 20, 000 x g for 20 min at 4°C to remove the membrane fraction. The supernatant was collected as soluble tau fraction [low-speed supernatant (LSS)] and stored at -80°C until further processed through Western blot. For the sarkosyl insoluble tau fraction, equal amounts of protein from the LSS were mixed with 10% sarkosyl solution to a final 1% sarkosyl concentration and incubated on a rotator for 30 min at room temperature. The samples were then centrifuged at 100, 000 x *g* in a Beckman TL-100 ultracentrifuge at 20°C for 1 h. The pellet was resuspended in 100 μl 1% sarkosyl solution and spun again at 100, 000 x g at 20°C for 1 h. The supernatant was then discarded, and the pellet air-dried for 30 min. Subsequently, 50 μl of modified O+ buffer (62.5 mM Tris-HCl, 10% glycerol, 5% β-mercaptoethanol, 2.3% SDS, 1 mM EDTA, 1 mM ethylene glycol-bis(β-aminoethyl ether)- tetraacetic acid (EGTA), 1 mM NaF, 1 mM Na_3_VO_4_, 1 nM PMSF and 1× cOmplete protease inhibitor cocktail, including about 1 μg/ml of bromophenol blue) was added and the sample vortexed for 1 min, and then boiled for 5 min [Sarkosyl Pellet (SP) fraction] and kept frozen at −80°C until used for Westerns. The LSS fraction (500 μg—about 30 μl) was diluted with the modified RIPA buffer (about 220 μl) and modified O+ buffer (50 μl), resulting in a final protein concentration of 25 μg/15 μl, and processed in the same way (boiled for 5 min and frozen). For Western blots, the soluble and insoluble tau fractions were thawed, reboiled for 5 min and electrophoresed (15 μl per lane) on a 12% (w/v) polyacrylamide gel. The proteins were then transferred to a nitrocellulose membrane that was subsequently blocked in 5% nonfat milk with 0.1% Tween-20 in TBS for 1 h, and incubated with different antibodies at 4°C overnight (Tau-5 (Santa Cruz, sc-58860, 1:1, 000, tau epitope 210–241), CP27 (1:1000, recognizes human tau epitope 130-150 but not mouse tau, generously provided as cell culture supernatant by Peter Davies) PHF1 (1:1, 000, tau epitope around aa P-Ser396, generously provided as cell culture supernatant by Peter Davies). Following washes, the membranes were then incubated for 2 h with 1:2, 000 horseradish-peroxidase (HRP) conjugated goat anti-mouse antibody (ThermoFisher Scientific), developed in ECL (ThermoFisher Scientific), imaged with Fuji LAS-4000, and the signal quantified with ImageQuant software. All the samples within each figure panel were run on the same gel, and each set of blots was repeated at least twice with one set used for quantitation.

### Acute tau antibody treatment

For acute immunization using the 5G2 antibody or the IgG1 isotype control, ThyG-1^CaMP6^: PS19 mice were first imaged via a window into the brain and then infused twice (100 µg each time) into the femoral vein under isoflurane anesthesia (3% induction, 1.5–2.0% maintenance), 4 days apart, before being imaged again on day 8. Cx3cr1^GFP^: PS19 mice and Cx3cr1^CreER^: PS19 mice were only imaged after 5G2 intravenous infusion.

### Microdialysis – tau antibody target engagement

The procedure began with a survival stereotaxic surgery on 7-8 month old female JNPL3 mice utilizing aseptic technique. The mice were anesthetized with isoflurane (3% induction, 1.5–2.0% maintenance) and placed in a stereotaxic frame on top of a heated pad to maintain a body temperature of ∼37 °C. The depth of anesthesia was monitored frequently during the surgery. An incision was made on the scalp exposing landmarks (bregma, lamda, and midline) to obtain the right injection co-ordinates. The skull was drilled at that location and a guide cannula (Amuza) was inserted into their left hippocampus (3.0 mm post bregma, +2.5 lateral, and 1.2 ventral from dura at a 12° angle) and secured via dental cement. Following a monitored recovery from surgery (30 min-1 h), a 2 mm probe (Amuza) was inserted into the guide cannula. The mice were then placed into the universal microdialysis cage (BASi) allowing them to move freely during collection of brain interstitial fluid (ISF). Sterile filtered artificial cerebral spinal fluid (aCSF) with 2% BSA (Sigma) was perfused through the probe at a rate of 1.2 μL/min with samples collected at 1.0 μL/min (1 hour intervals). Brain ISF was collected continuously for 48 h (day 1 as recovery, day 2 before and after treatment). Treatment animals received 5G2 tau antibody immunotherapy via reverse dialysis (n=5, 2 h infusion) following the establishment of a stable 24 h baseline period. Control mice (n=11) received only aCSF and underwent dialysate collection for 48 h uninterrupted. All infusion conditions were controlled across treatments. Sample analysis began at hour 16 and ended at hour 48. Animals were given a subcutaneous injection of analgesic buprenorphine (0.05-0.1 mg/kg) twice daily. Samples were quantified by an ELISA kit for total human tau (ThermoFisher Cat# KHB0041) as per manufacturer’s protocol to determine brain ISF tau levels before, during, and following treatment.

### Immunohistochemistry

Cx3cr1^CreER^ and Cx3cr1^GFP^ mice were deeply anesthetized with a mixture of ketamine (100 mg/kg) and xylazine (15 mg/kg) and then perfused with 10 ml phosphate buffered saline (PBS). Half of the brain was stored for Western blot at -80 C and the other half was immersed in 4% paraformaldehyde (PFA) in PBS at 4 degrees overnight. After PFA fixation, brains were rinsed with PBS and 100 µm sections were prepared with a vibratome (Leica VT1000S). Coronal sections from each brain were permeabilized in a buffer containing 1% Triton X-100 and 5% bovine serum albumin (BSA) in PBS for 3 hours. Sections were incubated overnight with primary antibodies against Iba-1 (Wako 1:400, rabbit) and PHF-1 (1:1000, mouse) in a buffer containing 0.1% Triton X-100 and 5% BSA. Sections were then washed 3 times with 0.05% Tween-20 in PBS and incubated overnight with Alexa Fluor-conjugated goat anti-rabbit antibodies (Invitrogen Cat. No. A11008, 1:500) in a buffer containing 0.1% Triton X-100 and 5% BSA. Sections were then washed 3 times with 0.05% Tween-20 and mounted for imaging. Confocal images of these coronal sections were obtained using a Zeiss 800 confocal microscope equipped with a 20X lens (N.A. 0.75) with a step size of 5 µm and a 63X lens (N. A. 1.4) with a step size of 0.5 µm. We generated a Z projection to create a single image for data analysis. Image processing and data acquisition of microglia morphology and Iba-1 expression were done using Fiji (version 2.14.0).

### Surgical preparation for imaging awake, head-restrained mice

These experiments were performed similarly to as we have described previously for a different tau antibody (Wu et al., 2020) except that instead of expressing the calcium indicator GCaMP6s via AAV in JNPL3 tauopathy mice, a transgenic cross was generated between PS19 P301S tau (JAX #008169) mice and a transgenic model, generously provided by Wen-Biao Gan at NYU, that has GCaMP6 expressed in neurons under the Thy-1 promotor (Cichon et al., 2019). The PS19 model is in many ways comparable to the JNPL3 model and is better suited to cross with the Thy-1^GCaMP6^, Cx3cr1^GFP^, or Cx3cr1^CreER^: tdTomato^flox^ mouse models because they share C57BL/6 strain background.

Briefly, surgery was performed using aseptic techniques under a mixture of ketamine (100 mg/kg) and xylazine (15 mg/kg) with the mouse remaining on a heated pad while anesthetized to maintain a body temperature of ∼37 °C. With the mouse in a stereotaxic frame, the skull was exposed via a midline scalp incision to reveal the landmarks (bregma, lambda, and midline) for the cranial window. A skull region (∼0.2 mm in diameter) was located over the primary motor cortex (based on stereotactic coordinates at 0.2 mm anterior to bregma and 1.2 mm lateral to midline) and marked with a pen to be carefully removed and replaced by a precut #1 square cover glass (World Precision Instruments, coverslip No. 1). Cyanoacrylate-based glue and dental cement were used to seal the edges of the cover glass, followed by skin suturing. The mouse was then placed on a heating pad and when awoken into its home cage.

For implanting the head holder, the mouse was anesthetized as indicated above, and its head shaved. The skull surface was then exposed with a midline scalp incision. The periosteum tissue over the skull surface was removed without damaging the temporal and occipital muscles. Two parallel micro-metal bars were attached to the animal’s skull to serve as the head holder to help restrain the animal’s head and reduce motion-induced artifacts during imaging. A skull region was located over the primary motor cortex and pen marked as described above. A thin layer of cyanoacrylate-based glue was first applied to the top of the entire skull surface. Then the head holder was mounted with dental acrylic cement (Lang Dental Manufacturing Co., IL. USA) such that the marked skull region was exposed between the two bars. The marked region for imaging was kept exposed. Next, we created the cranial window over the marked region. A high-speed drill was used to carefully reduce the skull thickness under a dissecting microscope (skull thickness ∼20 µm). The skull was immersed in artificial cerebrospinal fluid (aCSF) during drilling. For open skull preparation, a small, square craniotomy (1-1-5 mm diameter) was made and covered with a coverslip (World Precision Instruments, coverslip No. 1) fitting the size of the bone removed. The coverslip was glued to the skull to reduce the motion of the exposed brain. Before imaging, the mouse was given one day to recover from the surgery-related anesthesia and habituated for a few times (10 min each) in the imaging apparatus to minimize potential stress during head restraining and awake imaging. For a mouse that was implanted with a chronic glass window followed by three to four weeks of recovery, the head holder was implanted as described, and the glass window was cleaned before imaging. Once the implant was done, the mouse was put back into its home cage for recovery (Bai et al., 2017;Cichon and Gan, 2015;Li et al., 2017;Wu et al., 2020). A mouse implanted with a chronic window was housed individually throughout the study. When the window had cleared about four weeks later, the mouse underwent baseline two-photon imaging followed by re-imaging after the acute antibody treatment.

Analogous to our prior efficacious acute in vivo treatment studies with other tau antibodies (Congdon et al., 2016;Wu et al., 2018;Wu et al., 2020), the 5G2 tau antibody targeting Asp421 or its IgG1 control were injected (100 µg each time) into the femoral vein of the Thy-1^GCaMP6^: PS19, or Cx3cr1^CreER^: tdTomato^flox^ : PS19 mice on Day 1 after the baseline imaging and on Day 4, followed by reimaging on day 8.

The free-floating treadmill (101 cm × 58 cm × 44 cm) allowed head-fixed mice to move their forelimbs freely to perform motor running tasks. To minimize motion artifacts, it was constructed with the moving parts (motor, belt and drive shaft) isolated from the stage of the microscope and the supporting air-table. All the mice had intact motor function and could successfully run on the treadmill. PS19 mice develop motor impairment with age but it is not typically seen at the age of these mice.

Two-photon imaging was collected with an Olympus Fluoview 1000 two-photon system (920 nm) equipped with a Ti:Sapphire laser (MaiTai DeepSee, Spectra Physics). Ca^2+^ signals were recorded at 2 Hz for recording neuronal activity and 1Hz for detecting microglia, using a 25X objective (NA 1.05, 512 × 512 pixels) (Optical zooms 1.5 for L2/3 somas and 2 for microglia). The mice were forced to run (referred to as running) at the speed of 1.67 cm/s or allowed to rest (referred to as resting) on a treadmill with the head fixed on top of the custom-built free-floating treadmill. The purpose is to increase neuronal activity, which we have shown to facilitate detection of neuronal dysfunction in tauopathy mice as revealed by calcium imaging (Wu et al., 2020). Two-photon Ca^2+^ images were collected for five trials at 100 s each at different focal planes from GCaMP6-positive neurons within the motor cortex (150 - 300 µm below the pial surface) under resting or running conditions, performed within the same testing session. The same cells were imaged during these two conditions. For before vs after treatment, the same focal planes were imaged but because these images were collected on different days, there may have been a slight shift in the plane so the same cells may not have been analyzed as reflected in the statistical analysis (unpaired comparison).

The use of GCaMP6s and a 2 Hz sampling rate does not capture certain physiological responses such as patterns of neuronal action potentials. We chose to use the slow GCaMP6s, instead of the fast GCaMP6f, because it is more sensitive for detecting small changes in Ca^2+^ activity. The 2 Hz sampling rate was chosen to match the slow kinetics of GCaMP6s and to minimize the potential phototoxicity. GCaMP6s and 2 Hz sampling rate were used as previously reported (Bai et al., 2017;Cichon and Gan, 2015), and this particular Ca^2+^ indicator has been used for Ca^2+^ imaging in tauopathy mice by us and others (Marinkovic et al., 2019;Wu et al., 2020).

### Image analysis

Neuronal Ca^2+^ activity, indicated by GCaMP6 fluorescence changes, and microglial morphology, indicated by GFP or tdTomato expression, were analyzed using Image J software (NIH) as we have described previously (Wu et al., 2020). The GCaMP6 fluorescence (F) during resting and running was measured by averaging pixels within each soma of GCaMP6 positive neurons. The morphology of microglia, frequency of the Ca^2+^ transients, amplitude of the peak Ca^2+^ transient, and total Ca^2+^ activity (area under the curve, AUC) (Bruni et al., 2017;Overk et al., 2015;Reznichenko et al., 2012;Schmunk et al., 2017) were analyzed by using Graph Pad Prism V. 10.

All imaging stacks (microglia) and time-lapse images (L2/3 neurons) from each field of view were motion corrected using TurboReg plug-in for ImageJ (Thevenaz et al., 1998). Regions of interest (ROIs) corresponding to visually identifiable GCaMP6-expressing somas were selected manually. We excluded the cells that were active during resting conditions and chose those that displayed increased calcium activity upon running activation. The fluorescence time course of each whole field of view taken from the *Thy1*^GCaMP6^ expressing mice was measured by averaging all pixels within the ROIs. The *ΔF/F_0_* value was calculated as *ΔF/F_0_ = (F-F_0_)/F_0_*, in which *F_0_* is the baseline fluorescence signal averaged over ten seconds after background extraction. F0 and threshold were determined as the mean and three times the SD of the F values within the baseline interval.. The frequency of Ca^2+^ transients was calculated as the number of Ca^2+^ transients per 100 s for each soma. The amplitude of the peak Ca^2+^ transient was the highest amplitude value of the Ca^2+^ transients. The total Ca^2+^ activity (area under the curve, AUC) was the average of all the integrals over the time periods (100 s) above the threshold.

### Statistics

Statistical analyses were performed using GraphPad Prism (version 9.0). All data are reported as mean ± SEM. A value of P < 0.05 was considered statistically significant. Specific tests are described in the figure legend of each figure.

## RESULTS

### Antibody characterization

The two monoclonal antibodies (mAbs), 5G2 and 1G11, were selected from a group of monoclonals that specifically recognized the immunogen peptide tau 407-421 over a longer peptide tau 407-423, based on analysis of the hybridoma culture supernatants (data not shown). Of the two, 5G2 had much higher affinity for Tau-Asp421 than 1G11 based on Biacore analysis (**Figure 1A**, 5G2 K_D_ = 7.46 x 10^-9^ M vs. 1G11 K_D_ = 1.07 x 10^-6^ M). Furthermore, 5G2 had some binding to the longer peptide (K_D_ = 5.07 x 10^-6^ M), whereas 1G11 did not bind to it. Hence, higher affinity was associated with less specificity. Subsequently, immunohistochemistry of brain sections from an AD patient and a JNPL3 mouse revealed that both antibodies recognize tau pathology, with 5G2 resulting in stronger staining than 1G11, with no staining detected in control human brain or wild-type mouse brain (**Figure 1B-M**).

**Figure 1.**
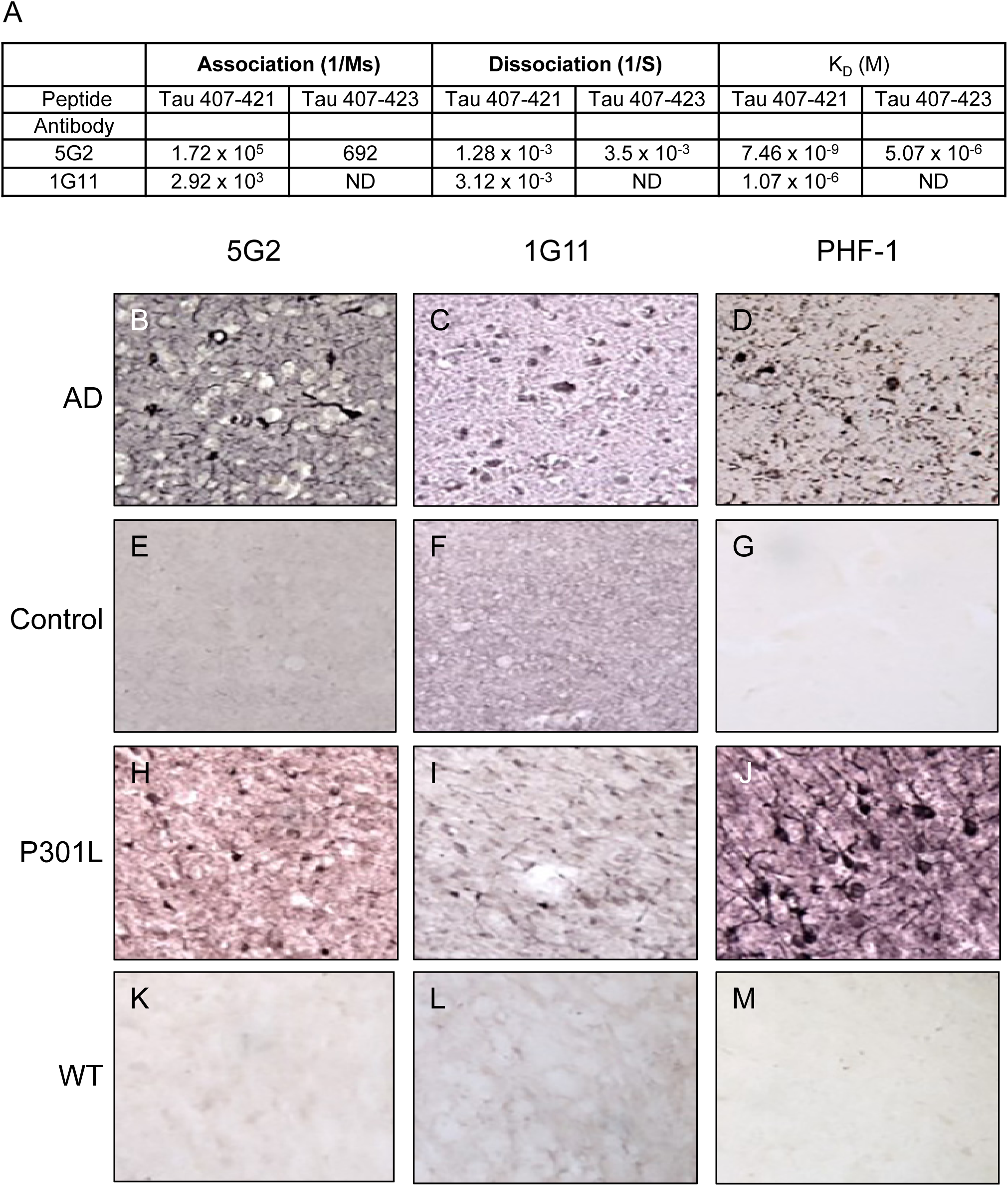
Characterization of the tau antibodies targeting truncated Asp421. **A)** Biacore assay demonstrated higher binding affinity for 5G2 (K_D_ = 7.46 x 10^-9^ M) compared to 1G11 (K_D_ = 1.07 x 10^-6^ M) for the truncated Asp407-421 peptide. The 5G2 showed some reactivity towards the control Asp407-423 peptide (K_D_ = 5.07 x 10^-6^), whereas the 1G11 antibody did not bind to it (ND: not detected). **B-D)** The 5G2 antibody showed greater reactivity towards tau pathology in Alzheimer’s brain tissue than the 1G11 antibody, which fits with its greater affinity for the Asp421 epitope. PHF-1 was used as a positive control, resulting in comparable staining of tau pathology as for the 5G2 antibody. **E-G)** Control human brain sections did not stain with any of the three tau antibodies, indicating their selectivity for pathological tau aggregates in the fixed brain tissue. **H-M**) Comparable pattern and intensity of staining was seen with these antibodies in JNPL3 tauopathy mouse brains, and lack of staining in wild-type brains.

### 5G2 treatment prevents PHF neurotoxicity and PHF-induced increase in total and phosphorylated tau in both primary tauopathy neurons and in mixed tauopathy cortical cultures

To promote tau pathology and to examine antibody-mediated clearance of tau and prevention of its toxicity, primary tauopathy neurons and mixed tauopathy cortical cultures were treated with PHF-enriched pathological tau protein (10 µg/ml) isolated from a human tauopathy brain, with or without Asp421 antibodies (5G2 and 1G11) each at 10 µg/ml for a duration of 24 h, 48 h, 72 h and 96 h. Western blots were conducted for NeuN, total tau, and phosphorylated tau at Ser199 (pSer199), to observe the effects of antibody treatment on PHF neurotoxicity and PHF-induced increase in total and phospho-tau protein.

### Primary neurons

Two-way ANOVA analyses of NeuN, total tau and pSer199 tau after treatment for 24-96 h showed significant overall effects for all three measurements (treatment: p < 0.0001; time: p < 0.0001; interaction: p < 0.0001). Neurotoxic effect of PHF administration was evident from 48 h through 96 h by reduced NeuN levels for the neurons treated with PHF alone (**Figure 2A, B**, 48 h: 22%, 72 h: 36% and 96 h: 49%, p < 0.0001 for all). Co-treatment with IgG was unable to prevent PHF toxicity at any of the time points. Treatment with the higher affinity antibody, 5G2, completely prevented PHF-induced neurotoxicity at all the time points (48-96 h, p < 0.0001 for all), resulting in NeuN levels comparable to untreated controls. The lower affinity antibody, 1G11, fully prevented toxicity at 48 h (p < 0.0001), and partially at 72 h and 96 h (21% and 44% reduction in NeuN compared to untreated control; p < 0.0001).

**Figure 2.**
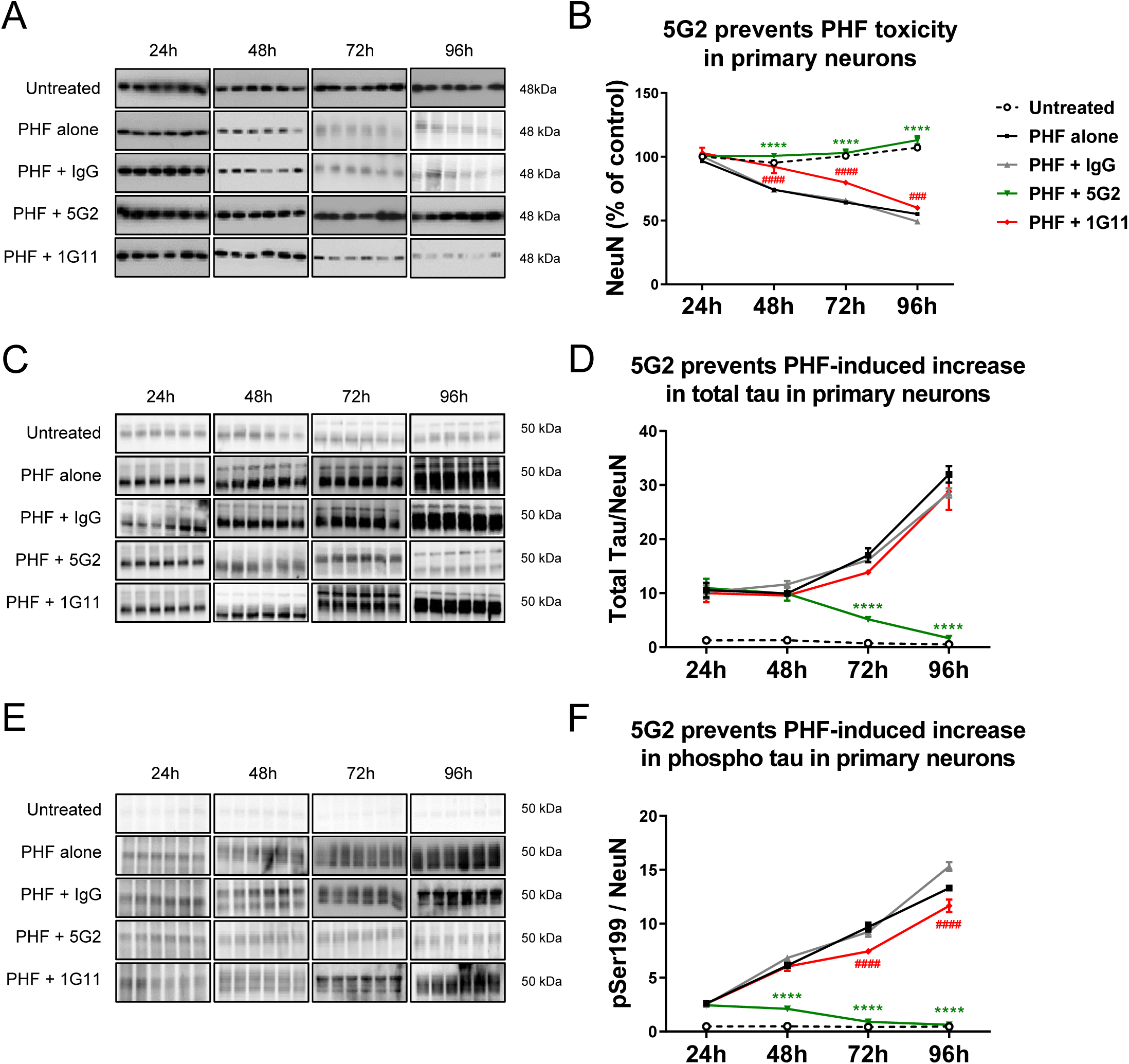
5G2 treatment prevented PHF-tau induced neurotoxicity and cleared total tau and phospho-tau in primary neuronal culture. Primary neurons from Day 0 P301L mice were co-treated with PHF (10 µg/ml) and Asp421 antibodies (5G2 or 1G11; 10 µg/ml) for a duration of 24 h, 48 h, 72 h and 96 h. Western blots were performed against NeuN, total tau and pSer199. Total tau and phospho-tau levels were normalized against NeuN. **A, B)** Quantification of NeuN levels demonstrated a significant PHF-induced neurotoxicity in the neurons treated with PHF alone and PHF + IgG controls with significant reduction in NeuN levels at 48 h (PHF alone: 22%, PHF + IgG: 23%), 72 h (PHF alone: 36%, PHF + IgG: 35%) and 96 h (PHF alone: 49%, PHF + IgG: 54%), when compared to the untreated cells, p < 0.0001 for all. 5G2 blocked this neurotoxicity resulting in NeuN levels comparable to that of the untreated cells. 1G11 was less effective but fully blocked PHF neurotoxicity at 48 h and partially at 72 h and 96 h (21% and 44% reduction in NeuN, respectively, p < 0.0001), compared to untreated controls. **C, D)** 5G2 effectively reduced total tau levels at 72 h (70%) and 96 h (95%), compared to PHF alone (p < 0.0001 for both). 1G11 and IgG were ineffective in reducing total tau levels at all-time points. **E, F)** A robust increase in pSer199 level was observed in all PHF-treated groups. 5G2 co-treatment gradually reduced pSer199 levels at 48 h (65%), 72 h (91%) and 96 h (95%) compared to the PHF alone group. 1G11 was less effective causing a subtle reduction in pSer199 level at 72h (19%) and 96h (12%) when compared to PHF alone (p < 0.0001 for both). IgG treatment was ineffective at all-time points. Statistical significance was determined by two-way ANOVA followed by Bonferroni post hoc test for multiple comparisons, n = 6, values are represented as mean + SEM. **** p < 0.0001 PHF + 5G2 vs. PHF alone or PHF + IgG, #### p < 0.0001 PHF + 1G11 vs. PHF alone or PHF + IgG, ### p < 0.001 PHF + 1G11 vs. PHF + IgG.

PHF addition to the culture increased total tau levels several fold compared to untreated neurons (**Figure 2C, D**). The 5G2 antibody decreased total tau levels at 72 h (70%, p < 0.0001) and 96 h (95%, p < 0.0001), compared to the neurons treated with PHF alone for the respective time points (**Figure 2C, D**). Control IgG or 1G11 were ineffective in clearing tau at all the time points.

Likewise, PHF addition increased phospho-tau levels several fold compared to untreated controls. The 5G2 antibody decreased phosphorylated tau levels at 48-96 h (**Figure 2E, F**, 48 h: 65%, 72 h: 91% and 96 h: 95% compared to PHF alone, p < 0.0001 for all). 1G11 treatment also proved effective in reducing pSer199 levels at 72 h (19%, p < 0.0001) and 96 h (12%, p < 0.0001), compared to cells treated with PHF alone for the respective time points. As for total tau, control IgG was ineffective in clearing phospho-tau at all the time points.

### Mixed culture

Two-way ANOVA analyses of NeuN, total tau, pSer199 tau, and Iba1 after treatment for 24-96 h showed significant overall effects for all four measurements (treatment: p < 0.0001; time: p < 0.0001; interaction: p < 0.0001). As in the primary neurons, PHF was neurotoxic in the mixed culture as reflected by decrease in NeuN levels in culture treated with PHF alone compared to untreated cells from 48 h through 96 h (**Supplementary Figure 1A, B**, 48 h: 24%, 72 h: 37% and 96 h: 54%, p < 0.0001 for all). This neurotoxicity was completely prevented by the 5G2 antibody for all the time points (p < 0.0001), with a significant increase observed in NeuN level at 96 h (21%, p < 0.0001) when compared to the untreated cells, thereby indicating neurotrophic effect of 5G2 treatment in this type of culture. The lower affinity antibody 1G11 partially inhibited PHF-induced neurotoxicity as reflected by reduction in NeuN levels by 20% and 26% at 72 h and 96 h respectively, in comparison to the untreated cells (p < 0.001-0.0001).

As in the primary neurons, total tau levels were increased several fold in the PHF-treated mixed culture, compared to untreated controls. The 5G2 antibody decreased total tau at 48-96 h compared to the culture treated with PHF alone (**Supplementary Figure 1C, D**, 48 h: 34%, 72 h: 39% and 96 h: 68%, p < 0.0001 for all). The 1G11 antibody and control IgG were ineffective at all time points except at 96 h where they reduced total tau compared to PHF alone group (18-19% reduction, p < 0.0001).

Likewise, phospho-tau levels were increased several fold in the PHF-treated culture, and these levels were robustly reduced with 5G2, compared to culture treated with PHF alone (**Supplementary Figure 1E, F**, 48 h: 74%, 72 h: 92%, 96 h: 96%, p < 0.0001 for all). 1G11 was ineffective in clearing phospho-tau at all time points except at 96 h (25% reduction compared to PHF alone, p < 0.0001).

Overall, these results in primary and mixed culture show strong efficacy of the higher affinity antibody, 5G2 in preventing PHF-mediated toxicity and in clearing total and phospho-tau in primary and mixed culture, whereas the lower affinity antibody, 1G11 is only partially effective.

### 5G2 treatment completely inhibits PHF-induced microglial activation

Microglial activation is associated with Alzheimer’s disease and microglia are likely to be involved in extracellular clearance of tau-antibody complexes. To determine the effects of Asp421 antibody treatment on microglial activation, Western blots were undertaken against Iba1, which is a marker for both resting and active microglia, using the mixed cortical cultures co-treated with PHF and Asp421 antibodies at 24 h, 48 h, 72 h and 96 h (**Supplementary Figure 1G, H**). This marker increases with microglial activation (Ito et al., 1998). Two-way ANOVA analysis revealed a highly significant treatment and time effect and interaction between the two (p < 0.0001 for all). PHF treatment led to time dependent increase in microglial activation as measured by Iba1 levels (98% - 1074% above untreated controls at 24 – 96 h, p < 0.01-0.0001). Similar increase was observed in the PHF + IgG control group (74% - 1072% above controls, p < 0.01-0.0001), and the lower affinity antibody, 1G11, had limited effect on attenuating this activation (54% - 855% above untreated control at 24 - 96 h, p<0.01-0.0001). In contrast, the high affinity antibody, 5G2, prevented PHF-induced microglial activation (24 – 53% above untreated controls at 24 – 96 h, not significant). The 5G2-mediated prevention of PHF-induced microgliosis differed significantly from the PHF alone group at 48 h (52% reduction, p<0.01), 72 h (86% reduction, p<0.0001) and 96 h (87% reduction p<0.0001). The more modest effect of 1G11 and control IgG differed significantly from the PHF alone group at 72 h (IgG 25% reduction, 1G11 36% reduction, p<0.0001 for both) and 96 h (1G11 19% reduction, p<0.0001). In summary, PHF treatment of mixed cortical cultures is associated with microglial activation, which is largely prevented by 5G2 and slightly attenuated by 1G11. This apparently beneficial effect of 5G2 on microglia may be secondary to its prevention of PHF-mediated neurotoxicity as microglia are typically activated following a neurotoxic insult.

### TRIM21 expression relates to cellular retention of antibodies, and varyingly to their acute (24 h) vs longer-term (5 days) efficacy, depending on their overall effectiveness

TRIM21 (T21) is a cytosolic Fc receptor that was earlier demonstrated to participate in the clearance of antibody-tau protein complex in culture (McEwan et al., 2017) and recently in a mouse model (Mukadam et al., 2023). To determine if this pathway was involved in the Asp421 antibody-mediated clearance of pathological tau protein, we knocked down (KD) this receptor in SH-SY5Y cells. Two-way ANOVA revealed a clear difference in T21 expression in T21 KD cells compared to naïve cells (p<0.0001) but the antibody groups did not differ significantly nor was there an interaction between the two factors (**Supplementary Figure 2A, B**). Therefore, the antibody treatment did not influence T21 expression. Post-hoc analysis revealed significantly reduced expression of T21 in all three groups (controls: 44%, p < 0.0001; 5G2: 41%, p < 0.0001; 1G11: 53%, p < 0.0001). A further Western blot analysis clearly illustrated a reduction in 5G2 (60%, p < 0.0001) and 1G11 (32%, p = 0.069) cellular retention in the T21 KD cells in comparison to their respective naïve cells (Two-way ANOVA: treatment: p < 0.0001, T21 expression: p < 0.0001, treatment x T21 interaction: p = 0.0004, **Supplementary Figure 2C, D**). This confirms the involvement of T21 in cellular retention of Asp421 antibodies, wherein a reduced expression of T21 clearly diminishes retention of Asp421 antibodies in the SH-SY5Y cells.

Having successfully established that T21 expression and KD were not influenced by the tau antibodies, differentiated naïve and T21 KD SH-SY5Y cells were pre-treated with PHF (5 µg/ml) for 24 h, followed by treatment with antibodies, 5G2, 1G11 or 4E6 (5 µg/ml) for 24 h and 5 days. With 24 h antibody treatment, two-way ANOVA analysis revealed a significant treatment effect (p < 0.0001) and an interaction between treatment and T21 expression (p < 0.0001) but T21 expression did not significantly influence the outcome (p = 0.702). 5G2 reduced total tau by 36% (p < 0.0001) and 19% (p < 0.01), in naïve and T21 KD cells, respectively, compared to cells treated by PHF alone (**Supplementary Figure 3A, B**). Within this treatment period, 1G11 was ineffective in both cells groups. The 5G2-mediated tau clearance between naïve and T21 KD cells was not significantly different, which at face value would seem to indicate that the T21 pathway is not involved in antibody-mediated tau clearance under these conditions. However, 5G2-mediated tau clearance in the naïve vs the T21 KD group was decreased by 47% (36% vs 19%, p = 0.056), which is comparable to their difference in T21 expression (41%). Thus, although not quite significantly different, this would seem to indicate that T21 expression relates acutely to antibody-mediated clearance of tau. To investigate whether this phenomenon is associated with epitope specificity, we assessed the efficacy of another anti-tau antibody, 4E6 against pSer396/404, which we have previously shown to be effective in clearing tau in various culture and in vivo assays under these same conditions (Congdon et al., 2013;Congdon et al., 2016;Congdon et al., 2019;Gu et al., 2013;Wu et al., 2018;Wu et al., 2020). The Westerns confirmed results similar to those seen with 5G2 treatment, with 4E6 mediating a 42% and 24% reduction in naïve and T21 KD cells, respectively (for both p<0.0001), compared to cells treated by PHF alone (**Supplementary Figure 3A, B**). Here, 4E6-mediated tau clearance did differ significantly between naïve and T21 KD cells (p < 0.05), supporting some involvement of the T21 pathway in tau clearance for 4E6 under these conditions. As for the 5G2 groups, 4E6-mediated tau clearance in the naïve vs the T21 KD group was decreased by 43% (42% vs 24%, p < 0.05), which again is comparable to the degree of T21 KD in this model. Overall, these findings suggest that T21 expression is linked to the acute efficacy of tau antibodies.

Considering this acute influence of the T21 pathway on antibody-mediated clearance of tau after 24 h treatment, the effect of longer treatment was assessed (**Supplementary Figure 3C, D)**. Both the naïve and T21 KD cells pre-treated with PHF (5 µg/ml) for 24 h, followed by treatment with antibodies 5G2, 1G11 and 4E6 each at 5 µg/ml for further 5 days resulted in a similar pattern as with the 24 h antibody treatment. However, with the 5 day treatment there was limited difference between the cell types (naïve vs T21 KD cells), in the reduction of total tau. With 5 day antibody treatment, two-way ANOVA analysis revealed a significant treatment effect (p < 0.0001) and an interaction between treatment and T21 expression (p = 0.040) but like at the shorter time period, T21 expression did not significantly influence the outcome (p = 0.171). A more pronounced antibody-mediated reduction in total tau levels vs PHF alone treated group was observed at the 5 day interval compared to 24 h in the naïve (5G2: 60% decrease, p<0.001; 1G11: 20% decrease, p < 0.05; 4E6: 61%, p < 0.0001) and in T21 KD cells (5G2: 55% decrease, p < 0.0001, 1G11: 3% increase, n.s.; 4E6: 50% decrease, p < 0.0001). At this longer treatment period, antibody efficacy in the naïve vs. T21 KD cells did not differ significantly for 5G2 (8% difference; 60% vs 55% decrease), or 4E6 (18% difference; 61% vs 50% decrease), but it did for 1G11 (p < 0.05). These findings indicate limited involvement of the T21 pathway in antibody-mediated tau clearance under these longer-term treatment conditions. Specifically, it is not involved for the highly efficacious antibodies 5G2 and 4E6 but it is involved for the less efficacious antibody 1G11.

GAPDH was considered as an internal control for these set of experiments, wherein no significant alteration was observed in either the naïve or T21 KD cells treated with 5G2, 1G11 and 4E6 antibodies for both 24 h and 5 d treatment in comparison to their respective cells treated with PHF alone (**Supplemental Figure 4**).

### Chronic 5G2 immunization reduces insoluble brain tau levels in tauopathy mice

Considering the consistent efficacy of 5G2 antibody treatment in the three different culture models, a further study was undertaken wherein 7-8 month old JNPL3 mice were immunized with 5G2 to determine the effect of Asp421 antibody treatment on clearance of tau in vivo. The animals received weekly intraperitoneal injections of 10 mg/kg of 5G2 or control IgG for 13 weeks. Brain analysis at the end of the study revealed a reduction in the antibody treatment group in insoluble tau (**Figure 3A-D**, CP27, 59%, p = 0.004, Tau-5, 74%, p = 0.007, PHF-1, 69%, p = 0.015), and a trend for a decrease in soluble tau (**Figure 3E-H**, Tau-5, 35%, p = 0.09).

**Figure 3.**
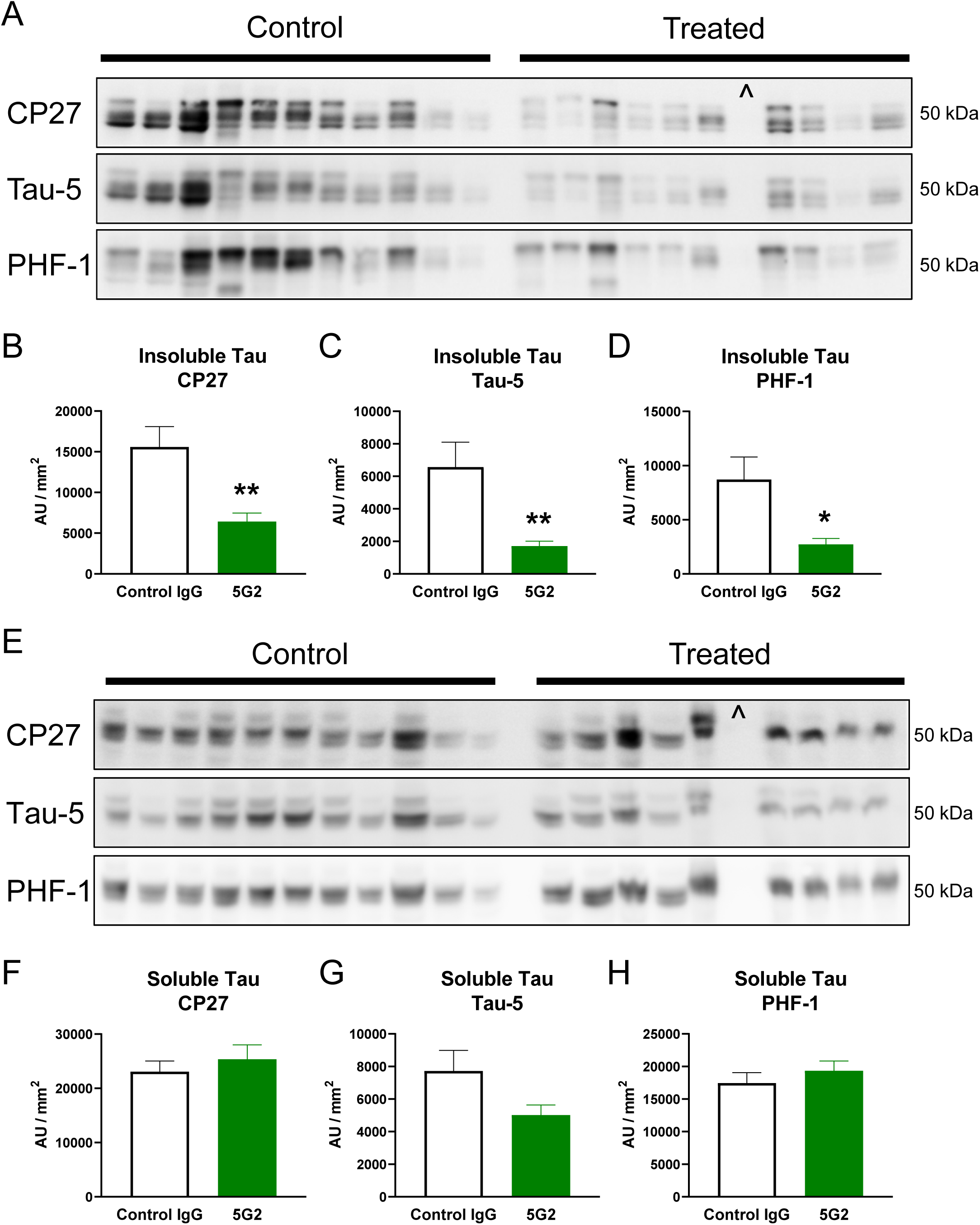
Chronic 5G2 immunization reduces insoluble brain tau in JNPL3 mice. The mice received 13 weekly 10 mg/kg intraperitoneal injections of 5G2 (n = 10) or control IgG (n = 11), followed by western blots of insoluble and soluble tau. **A-D)** Insoluble human tau, total tau and phospho-tau were reduced in the treated group by 59% (CP27, p = 0.004), 74% (Tau-5, p = 0.007) and 69% (PHF-1, p = 0.015). **E-H**) Soluble tau was not significantly reduced by the treatment although there was a strong trend for reduction in Tau-5 immunoreactive soluble tau (35%, p = 0.09). Statistical significance was determined by an unpaired t-test. Values are represented as mean + SEM. ^ marks an animal that did not express tau and was therefore excluded from the analysis.** p < 0.01, * p < 0.05.

### Acute 5G2 immunization reduces tau in brain interstitial fluid in tauopathy mice

A second in vivo study was conducted in 7-8 month old female JNPL3 mice, in which a microdialysis probe was implanted into their hippocampus. This allowed continuous sampling of their brain interstitial fluid (ISF) before, during and after treatment with a single dose of 5G2 (50 µg over 2 h), compared to control animals (vehicle aCSF). Tau levels in ISF show a diurnal fluctuation with the highest levels during the dark hours of the day (**Figure 4A**). The infusion of the antibody into the microdialysis probe prevented this increase throughout the 24 h sampling period following the start of the antibody treatment (**Figure 4A**, two-way ANOVA: treatment: p = 0.02, time: p = 0.03, treatment x time interaction: p < 0.0001). Specifically, during the dark period, 5G2 decreased tau levels by 46% (**Figure 4B**, t-test, p < 0.0001).

**Figure 4.**
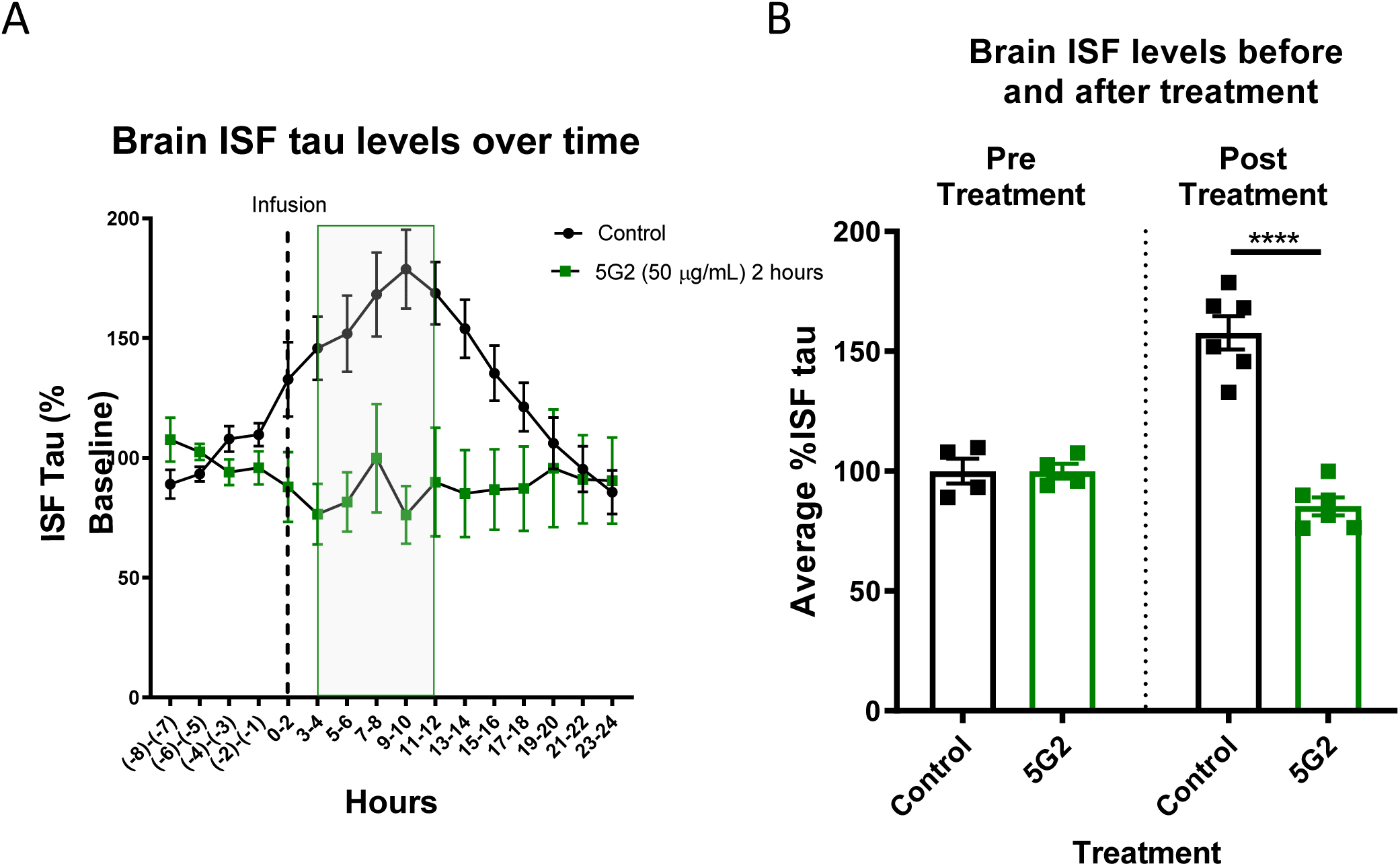
Acute 5G2 immunization reduces tau in brain interstitial fluid (ISF) in JNPL3 mice. **A)** Using in vivo microdialysis, a single infusion (50 µg/ml over 2 h dashed line in A) of 5G2 significantly reduced tau levels in brain ISF in 7-8 month old females, compared to controls (two-way ANOVA: treatment: p = 0.02, time: p = 0.03, treatment x time interaction: p < 0.0001). Note that tau levels in controls increase diurnally, peaking during the night (shaded area). **B)** Comparing pre- and post-treatment (shaded area in A) averages more clearly shows treatment-induced decrease of ISF tau levels by 46% (control: n = 11, 5G2 treatment: n=5). **** p < 0.0001.

### PS19 mice display altered calcium activity in L2/3 pyramidal neurons in the motor cortex

To determine how early neuronal deficits can be detected in tauopathy mice, we visualized calcium activity in neuronal soma at different ages during the progression of tau pathology (1 to 6 months old, n = 4-5 pairs per group/age, **Figure 5A**). We crossed the tauopathy mouse model PS19 with the Thy-1^GCaMP6^ mouse model (**Figure 5B**), which allowed us to assess calcium activity at layer 2/3 (L2/3) pyramidal neurons of the motor cortex.

**Figure 5.**
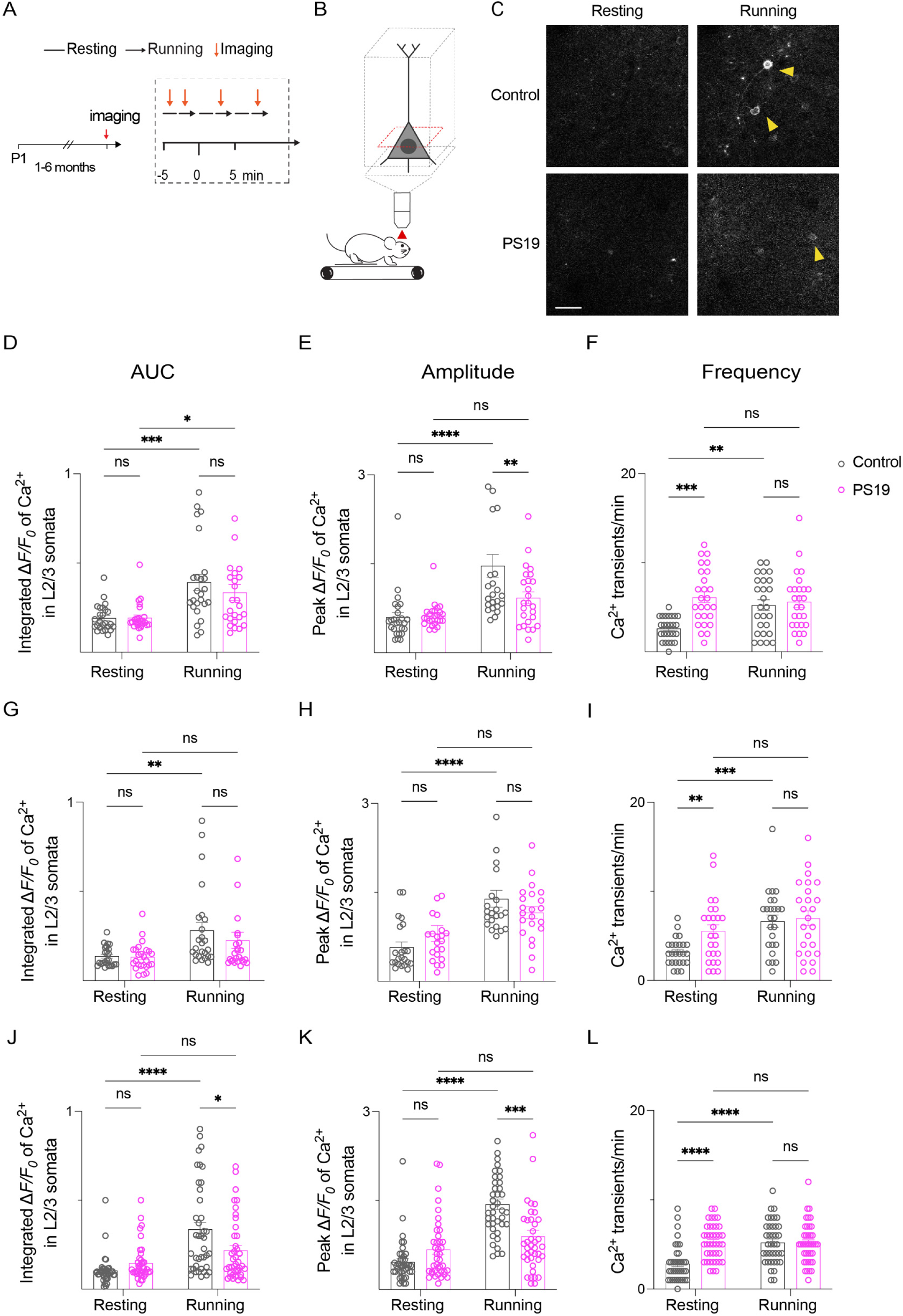
PS19 mice display altered calcium activity in L2/3 pyramidal neurons in motor cortex. **A, B)** Thy-1^GCaMP6^ mice were crossed with the PS19 mice. Data was collected at resting and running conditions. **C)** Two-photon images from control and PS19 mice showing calcium activity in a running state compared to the resting state. Scale bar: 20 µm. Mice of 1 to 6 months of age were used (n = 4-5 pairs per group/age). L2/3 pyramidal cells from control and PS19 mice of **D**-**F**) 1 to 2, **G-I**) 3 to 4, and **J-L**) 5 to 6 months old were recorded at resting and running periods to assess their total calcium activity (area under the curve, AUC), peak amplitude (amplitude), and frequency (number of calcium transients per 60 seconds). We selected cells that underwent increased calcium activity upon running activation. We averaged 20 to 30 cells from each age group/genotype tested. **D, G, J)** Total activity: two-way ANOVA: control resting vs control running: **D**, p = 0.0004; **G**, p = 0.0061; **J**, p < 0.0001). **D**) Total activity, two-way ANOVA: PS19 resting vs PS19 running: p = 0.0133). **E, H, K)** Peak amplitude: two-way ANOVA: control resting vs control running: **E**, p < 0.0001; **H**, p < 0.0001; **K**, p < 0.0001). **E, K)** Peak amplitude: two-way ANOVA: control running vs PS19 running: **E**, p = 0.0096; **K**, p = 0.0001. **F, I, L)** Frequency: two-way ANOVA: control resting vs control running: **F**, p = 0.0066; **I**, p = 0.0001; **L**, p < 0.0001; two-way ANOVA: control resting vs PS19 resting **F**, p = 0.0003; **I**, p = 0.0094; **L**, p < 0.0001). Multiple comparisons were conducted with Tukey’s post hoc test.

Total calcium activity (AUC) in control mice and PS19 mice from 1-2 (**Figure 5D-F**), 3-4 (**Figure 5G-I**), and 5-6 (**Figure 5J-L**) months of age were recorded at resting and running conditions. We detected higher total activity in L2/3 pyramidal neurons of control mice while running, compared to the resting state across all the age groups (**Figure 5D, G, J**, two-way ANOVA: control resting vs control running: **D**, p = 0.0004; **G**, p = 0.0061; **J**, p < 0.0001). In contrast, only the PS19 mice at 1-2 months of age increased their total activity while running vs resting (**Figure 5D**, two-way ANOVA: PS19 resting vs PS19 running: p = 0.0133). While running, control and PS19 mice did not differ in their neuronal calcium total activity except at 5-6 months of age (**Figure 5J**, two-way ANOVA: control running vs PS19 running: p = 0.0235).

Peak amplitude in control mice increased while running vs resting in all age groups (**Figure 5 E, H, K**, two-way ANOVA: control resting vs control running: **E**, p < 0.0001; **H**, p < 0.0001; **K**, p < 0.0001). In contrast, PS19 mice did not increase their peak amplitude upon running. While running, control mice had higher peak amplitude than PS19 mice at 1-2 and 5-6 months of age (**Figure E, K,** two-way ANOVA: control running vs PS19 running: **E**, p = 0.0096; **K**, p = 0.0001).

Regarding frequency of calcium transients, control mice increased their number of calcium transients upon running in all age groups (**Figure 5F, I, L**, two-way ANOVA: control resting vs control running: **F**, p = 0.0066; **I**, p = 0.0001; **L**, p < 0.0001). In sharp contrast, PS19 mice had comparable frequency of calcium transients under resting and running conditions. Their frequency was higher than in control mice in all age groups (**Figure 5F, I, L**, two-way ANOVA: control resting vs PS19 resting **F**, p = 0.0003; **I**, p = 0.0094; **L**, p < 0.0001). These data indicate that neuronal deficits occur before any overt detection of tau pathology.

### Acute 5G2 immunization improves neuronal function in tauopathy mice

Since the tauopathy mice had an abnormal neuronal Ca^2+^ profile, we examined if this could be corrected with acute tau antibody treatment (**Figure 6A-C**). Total calcium, peak amplitude, and frequency were analyzed in L2/3 somas from PS19 mice before and after (days 0 and 8) two intravenous doses of 5G2 antibody (n = 8) or IgG control (n = 5) on days 1 and 4 (100 µg each). Similar to the prior study, the Ca^2+^ profile of the untreated tauopathy mice did not differ between resting and running states at day 0 (**Figure 6D-F**). After 5G2 tau antibody treatment on day 8, all three Ca^2+^ parameters increased while the animals were running compared to resting (**Figure 6D-F**, two-way ANOVA: PS19 resting vs PS19 running **D**, p = 0.0002; **E**, p < 0.0001; **F**, p < 0.0001), analogous to what was observed in the non-tauopathy mice (**Figure 6J-L**). While running, both total calcium activity and amplitude were higher in 5G2-treated mice compared to the untreated PS19 mice (**Figure 6D, E**, two-way ANOVA: 5G2-treated PS19 running vs untreated PS19 running, **D**, p = 0.0010; **E**, p = 0.0001). Moreover, the frequency of Ca^2+^ transients in the 5G2-treated PS19 mice was lower than in untreated PS19 mice (**Figure 6F**, two-way ANOVA: 5G2-treated PS19 resting vs untreated PS19 resting, **F**, p = 0.0007). In contrast, control IgG1 was ineffective in improving neuronal function in PS19 mice (**Figure 6G-I**). Brain tissue analysis at the end of the study revealed a reduction in insoluble (CP27) and soluble (5G2) tau in the 5G2 antibody-treated group (**Figure 6J-L**, unpaired t-test: PS19 control IgG treated mice vs 5G2-treated PS19 mice, **K**, p= 0.0417; **L**, p = 0.0098). PHF-1 and CP27 reactive soluble tau did not differ between 5G2-treated and IgG control PS19 mice (**Supplementary Figure 5A-C**). Together, these results indicate that 5G2 treatment can restore neuronal activity to normal levels in tauopathy mice that is associated with clearance of insoluble tau and Asp421 truncated soluble tau.

**Figure 6.**
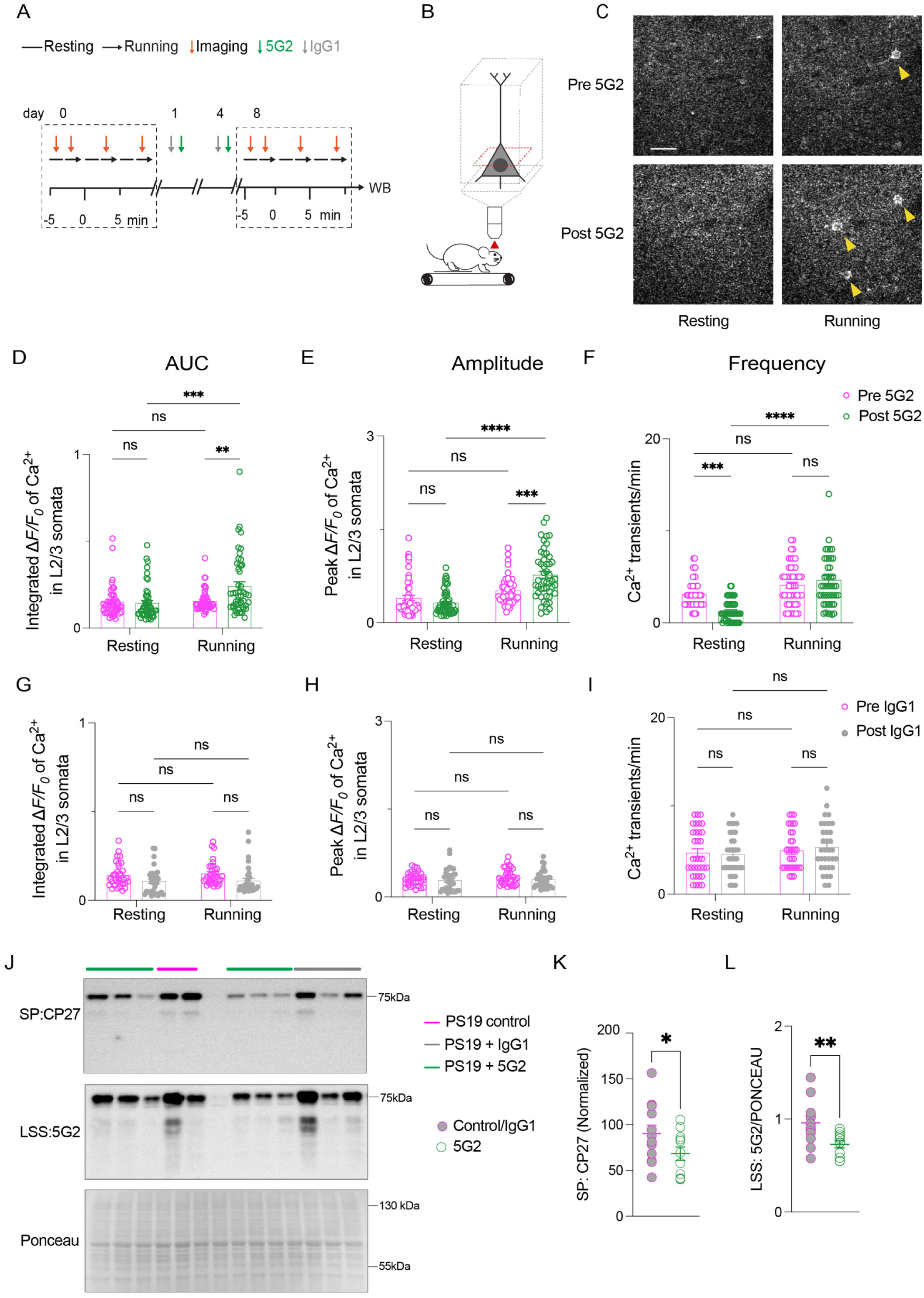
Acute 5G2 immunotherapy restores L2/3 neuronal function in PS19 mice. A, B) L2/3 somas from PS19 mice before and after two intravenous doses of 5G2 antibody (n = 50 somas from 8 mice) or IgG isotype control (n = 30 somas from 5 mice) on days 1 and 4 (100 µg each) were longitudinally imaged. **C)** Two-photon images from L2/3 pyramidal cells at resting and running conditions, before and after 5G2 antibody treatment. **D-F)** two-way ANOVA: PS19 resting vs PS19 running: **D**, p = 0.0002; **E**, p < 0.0001; **F**, p < 0.0001. **D, E)** two-way ANOVA: 5G2-treated PS19 running vs untreated PS19 running: **D**, p = 0.0010; **E**, p = 0.0001. **F**) two-way ANOVA: 5G2-treated PS19 resting vs untreated PS19 resting: **F**, p = 0.0007. **G-I)** IgG-treated mice showed no effect in any of the parameters tested. **J)** Representative blot showing CP27 and 5G2 signal in the insoluble and soluble fractions, respectively (n = 5 PS19 mice (control), n = 5 IgG-injected PS19 mice, n = 11 5G2-injected PS19^+/-^ mice. **K, L**) Unpaired T-test: PS19 control/IgG treated mice vs 5G2-treated PS19 mice, **K**, p = 0.0417, (CP27 signal); **L**, p = 0.0098 (5G2 signal). Multiple comparisons were conducted with Tukey’s post hoc test.

### Acute 5G2 tau immunotherapy reverses microgliosis in PS19 mice

We also examined whether anti-tau immunotherapy influences microglia structure in tauopathy mice. Cx3cr1^CreER^ mice were first crossed with flox-td-Tomato mice (control mice) and then with PS19 mice to obtain red microglia (n = 3 per genotype, 7-9 months old). The mice were then imaged after two intravenous doses of 5G2 antibody (100 μg each), 3 days apart (**Figure 7A-B**). Compared to control mice, PS19 mice had a larger microglia soma but 5G2 antibody treatment restored it to control size (**Figure 7C-D**, one-way ANOVA, control vs PS19 mice, **D**, p < 0.0001; PS19 mice vs 5G2-treated PS19 mice, p < 0.0001). We also assessed Iba-1 expression in the three groups and found increased expression of Iba-1 in PS19 mice compared to control mice but the 5G2 antibody treatment reduced Iba-1 levels to control values (**Figure 7E-F**, one-way ANOVA, control vs PS19 mice, **E**, p < 0.0001; PS19 mice vs 5G2-treated PS19 mice, p = 0.0014). Together, these results indicate that 5G2 treatment reverses microgliosis in tauopathy mice.

**Figure 7.**
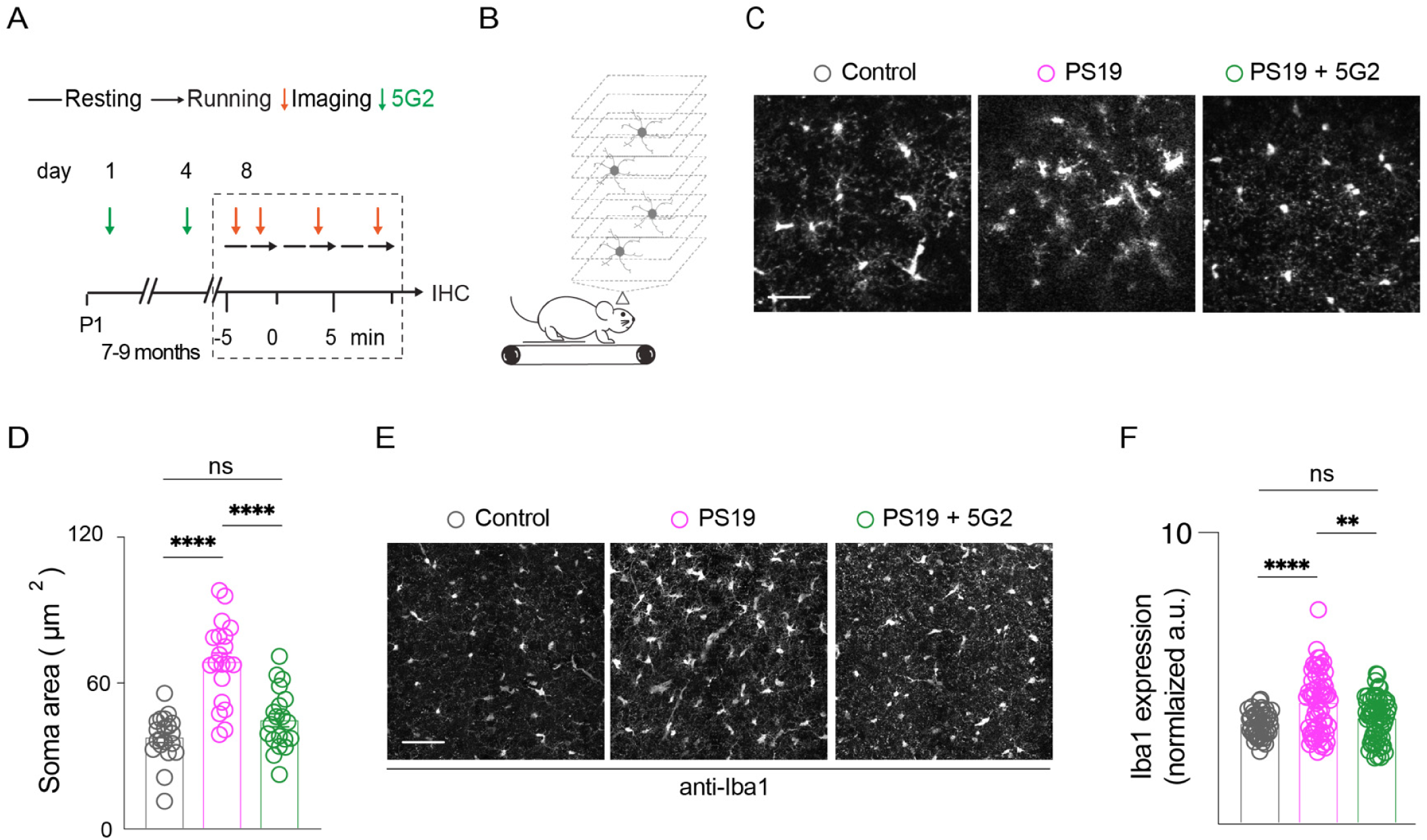
Acute 5G2 immunotherapy restores microglia morphology and Iba-1 expression in PS19 mice. **A, B)** Cx3cr1^CreER^: tdTomato^flox^ mice were crossed with the PS19 mice. 7 to 9-month-old control and PS19 mice (n = 3 per genotype) were imaged after administering two doses of 5G2 antibody. **C)** Two-photon images showing tdTomato expressing microglia. Scale bar: 10 µm. **D)** Ordinary One-way ANOVA, post hoc Sidak multiple comparisons: control vs PS19 mice, p < 0.0001; PS19 mice vs 5G2-treated PS19 mice, p < 0.0001 (n = 30 microglia cells per genotype). **E)** Two-photon images. Scale bar: 50 µm. **F)** Ordinary One-way ANOVA, post hoc Sidak multiple comparisons: control vs PS19mice, p < 0.0001; PS19 mice vs 5G2-treated PS19 mice, p < 0.0014 (n = 60 cells per genotype).

### Tau pathology precedes changes in soma size in microglia

We then sought to determine if structural changes in microglia precede the appearance of tau aggregates. Towards this aim, we crossed PS19 mice with Cx3cr1^GFP^ mice (control mice). Two to three-month-old control and PS19 mice were imaged (**Figure 8A-B**), with no differences seen between genotypes in the size of microglia soma (**Figure 8C-D**). At this age, tau pathology was readily detected in the PS19 mice as revealed by anti-phospho-tau staining (PHF1 antibody) in the motor cortex (**Figure 8E**) and CA3 area of the hippocampus (**Figure 8F**). These data indicate that tau pathology precedes changes in soma size of microglia.

**Figure 8.**
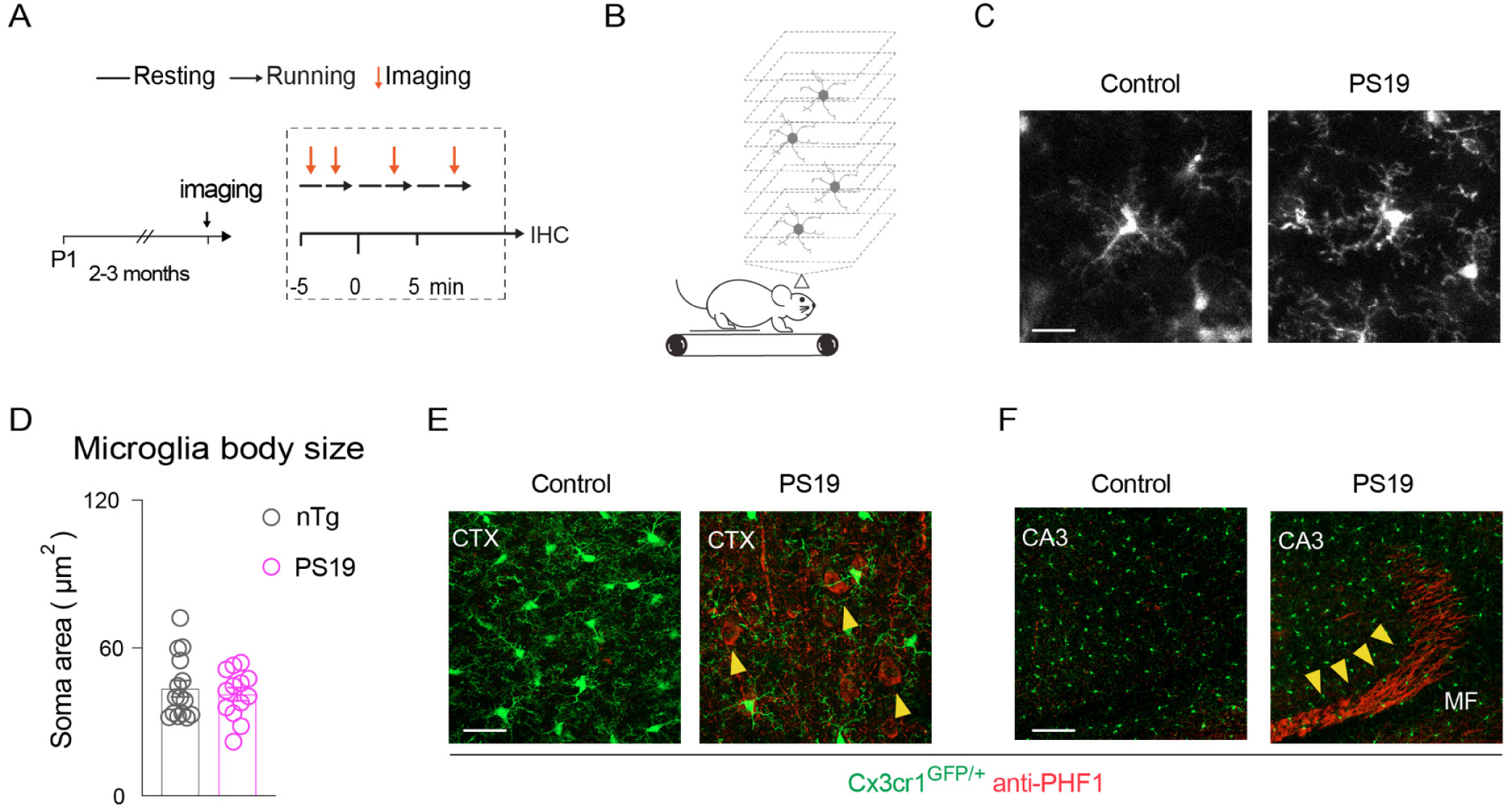
PS19 mice display tau accumulation in motor cortex and hippocampus without inducing changes in microglia soma size. **A, B)** Cx3cr1^GFP^ mice were crossed with the tauopathy mouse model PS19. Two to three-month-old control and PS19 mice (n = 4 per genotype) were 2P imaged. We examined microglia soma size and found no differences between genotypes. **C)** Two-photon images showing a microglia cell from control and PS19 mice. Scale bar: 20 µm. **D)** Quantification of microglia soma area (µm^2^) (n = 20 cells per genotype). **E, F)** Fixed brain slices from control and PS19 mice were incubated with the anti-tau antibody PHF-1. Confocal images were obtained to co-visualize microglia (green) and PHF-1-expressing cells (red) in the cortex (**E**, 60X, scale bar: 10 µm) and hippocampus (**F**, 10X, scale bar: 200 µm).

## DISCUSSION

Our study shows that targeting truncated tau at Asp421 with monoclonal antibody 5G2 prevents tau-mediated toxicity and leads to clearance of pathological tau both in primary tauopathy neurons, and in a mixed tauopathy cortical culture. In the latter model, this beneficial effect was associated with complete prevention of tau-mediated microgliosis, indicating an anti-inflammatory effect of the tau antibody treatment, presumably because of its prevention of neurotoxicity. The therapeutic benefits of the 5G2 antibody were evident as well in a human differentiated neuron-like model. Importantly, these beneficial effects were then confirmed in vivo, showing that 5G2: 1) cleared tau in brain interstitial fluid following a single dose as examined by microdialysis in awake JNPL3 tauopathy mice; 2) cleared insoluble tau in the same tauopathy mouse model following a chronic treatment, and; 3) improved neuronal function in PS19 tauopathy mice, as examined by calcium imaging, and cleared tau after an acute treatment. Mechanistically, the high affinity intracellular Fc-receptor and ubiquitin E3 ligase, TRIM21, was linked to intracellular retention of the tau antibodies. TRIM21 impacted antibody-mediated tau clearance at acute time points, but only to a limited extent long-term.

The benefits of targeting the truncated tau Asp421 epitope are comparable to our previous findings targeting the P-Ser396, 404 tau epitope (Asuni et al., 2007;Boutajangout et al., 2010b;Boutajangout et al., 2011;Congdon et al., 2013;Congdon et al., 2016;Congdon et al., 2019;Gu et al., 2013;Krishnamurthy et al., 2011;Krishnaswamy et al., 2020;Rajamohamedsait et al., 2017;Rosenqvist et al., 2018;Shamir et al., 2016;Shamir et al., 2020;Wu et al., 2018;Wu et al., 2020). However, here the higher affinity antibody was more effective (5G2 vs 1G11), whereas for the latter epitope, a lower affinity antibody was more efficacious (4E6 vs. 6B2). This is not particularly surprising. The Asp421 epitope has been linked to tau toxicity and seeding and these effects are likely neutralized by strongly capping it with an antibody. The P-Ser396, 404 epitope is known to be conformational with each phosphorylation site influencing the conformation of the other one. Our previous findings indicated that the effective tau antibody, 4E6, primarily bound to soluble pathological tau, whereas the ineffective antibody, 6B2, had a much higher affinity for insoluble tau, which may lead to neutralization of the antibody (Congdon et al., 2016). Furthermore, since 6B2 does not clear insoluble tau in tauopathy mice, high affinity to that particular epitope may render the aggregates more compact and less amenable to degradation (Congdon et al., 2016;Wu et al., 2018).

The high affinity cytosolic Fc receptor and ubiquitin E3 ligase, TRIM21 has previously been shown to be involved in tau antibody-mediated prevention of tau seeding (McEwan et al., 2017;Mukadam et al., 2023). Our findings indirectly support the binding phenomenon of this interaction and its acute but to a lesser extent its long-term involvement in antibody-mediated clearance of pathological tau. First, cellular retention of the tau antibodies was reduced by a comparable degree as the TRIM21 KD (about 50%), indicating their intracellular binding. Although the overall outcome of the two-way ANOVA analysis following the acute 24 h treatment indicated no significant effect of TRIM21 in antibody efficacy, a closer examination revealed it to be likely present under these acute conditions for the two efficacious antibodies, 5G2 and 4E6. Specifically, knocking down this receptor by about 50% in a human tauopathy neuron-like cellular model did reduce the acute (24 h treatment) but not the long-term (5-day treatment) efficacy of the highly effective tau antibodies 5G2 and 4E6 in clearing tau by a similar percentage. The modestly effective antibody, 1G11, only cleared tau in the naïve cells at 5 days but not in the TRIM21 KD cells at that time point. Overall, this TRIM21-antibody-mediated tau clearance appeared to be mostly transient or overtaken by presumably lysosomal-antibody-mediated tau clearance during the longer treatment period. We have previously shown that various tau antibodies are taken up into the endosomal-lysosomal system within neurons in which they bind to pathological tau (Congdon et al., 2013;Congdon et al., 2019;Gu et al., 2013;Krishnamurthy et al., 2011;Krishnaswamy et al., 2014;Shamir et al., 2016;Shamir et al., 2018;Shamir et al., 2020). We have hypothesized that antibodies against certain tau epitopes may loosen up the aggregates within the lysosomes and thereby facilitating their lysosomal clearance (Asuni et al., 2007). Another issue to consider is that TRIM21 has low brain expression (Uhlen et al., 2015;Zhang et al., 2014). Since it will be degraded alongside the antibody-target complex, its mediated tau clearance is likely to be only transient for highly efficacious antibodies, whereas it may take time to see its effect for less efficacious antibodies, as our findings indicate.

Regarding apparent differences between our findings and those by McEwan et al, our experimental design is not identical to that in their report that more closely links tau antibody efficacy to its binding to intracellular TRIM21. There, TRIM21 was completely knocked out, and the antibodies bound to different epitopes (tau19-46 and tau428-441). In addition, that prior study used HEK293 cells or undifferentiated SH-SY5Y cells and lipofectamine to enhance uptake of tau-antibody complexes. We have previously shown lack of tau antibody efficacy in an undifferentiated tauopathy SH-SY5Y model, whereas the same antibody was effective in clearing tau in an otherwise identical differentiated model (Shamir et al., 2020). Antibody uptake in the former undifferentiated model was exclusively via non-specific bulk uptake, whereas in the latter differentiated model, it was primarily via low affinity FcII/FcIII antibody uptake (Shamir et al., 2020). It is unclear if/how these different uptake mechanisms would influence antibody efficacy, and in both models we observed the antibodies within the endosomal/lysosomal system associated with the tau protein (Congdon et al., 2019;Shamir et al., 2016;Shamir et al., 2018;Shamir et al., 2020), as well as in neuronal and brain slice cultures and in vivo models (Congdon et al., 2013;Congdon et al., 2019;Gu et al., 2013;Krishnamurthy et al., 2011;Krishnaswamy et al., 2014). It is conceivable that the undifferentiated neuroblastoma cells rely more on exocytosis for protein clearance than their more neuron-like differentiated counterpart. Regardless of these discrepancies, our findings support antibody binding to cytosolic TRIM21 as reflected in a comparable reduction (about 50%) in intracellular antibodies as the degree of TRIM21 knockdown, and the involvement of this intracellular receptor primarily in acute but to a limited extent in long-term antibody-mediated tau clearance.

Microglia have emerged as central players in neurodegeneration but the cellular and molecular mechanisms behind their reactive behavior seen in this context are unclear. The enlargement of microglial soma toward an ameboid-like shape is a hallmark of its reactive phenotype, which is present in cells interacting with amyloid plaques and tau aggregates (Leng and Edison, 2021). We crossed the Cx3cr1^CreER^: tdTomato^flox^ mice with the PS19 mice to visualize microglia in vivo. As expected, we found larger somas and higher Iba-1 expression in microglia from PS19 mice of 7-9 months of age than in control mice, but the acute treatment of 5G2 in PS19 mice reverted microglia to a non-reactive control condition compared with the untreated PS19 mice. We also crossed the Cx3cr1^GFP^ mice with the PS19 mice to visualize microglia in mice at 2-3 months of age to examine if their reactive phenotype appears before or after the appearance of tau aggregates. While the soma size of microglia was comparable in control and PS19 mice, tau deposits were evident in both the CA3 region of the hippocampus and in cortex of the PS19 mice. These data, along with the abnormal neuronal calcium activity observed in PS19 mice at 1-2 months of age, strongly suggests that tau deposition and dysregulated neuronal activity precede microglia activation.

Microglia in culture are known to behave differently than microglia in vivo but our animal findings were mirrored in our culture studies. In our mixed tauopathy cortical model, microgliosis was clearly evident following PHF-tau addition to the culture, resulting in over 1000% increase in Iba1 levels. This inflammatory response was slightly attenuated by 1G11, the less effective antibody, and completely by 5G2, the highly effective antibody. Together with the in vivo findings, this bodes well for its therapeutic potential as it also relates well to its beneficial effects on neurons that were seen in the animals as well as in the primary neuronal and mixed cellular cultures .

Microgliosis is closely associated with amyloid-β plaques in AD and related models, where microglia infiltrate and encapsulate the plaques. It is less prominent in tau pathology, presumably because most of tau pathology is intracellular. In the mixed culture system, PHF enriched tau is added to the culture, in which it may stimulate microglial phagocytosis to an artificial degree either directly or indirectly via its neurotoxicity. However, it is a convenient system to examine acute inflammatory effects of pathological tau. In vivo, the situation is likely to be different. For example, we previously reported similar modest degree of microgliosis in htau/PS1 mice and related models with less tau pathology, thereby not clearly linking tau pathology to microgliosis in these mouse models (Boutajangout et al., 2010b). Likewise, we have reported that chronic tau immunotherapy does not affect the modest microgliosis observed in htau/PS1 or JNPL3 mice (Boutajangout et al., 2010b;Boutajangout et al., 2011). Because of the gradual chronic antibody-mediated clearance of tau pathology in these models, any potential effect on microglia is likely to have subsided at the end of the study. Outcome may be different in acute studies like in our mixed culture model, where the tau antibody clearly blocked microgliosis, directly or indirectly, and in the PS19 animals where a similar effect was seen. For a related insight into this issue, we previously examined tau antibody uptake into different cell types in cultured brain slices from tauopathy mice (Gu et al., 2013). In that model, about 80% of the intracellular tau antibodies were detected within neurons and about 10% within microglia with the remaining 10% not clearly associated with any particular cell type. This may be explained by greater antibody turnover in microglia because of their robust phagocytosis.

The 5G2 antibody was not only consistently effective in three different culture models but also in vivo during chronic or acute conditions. The chronic weekly 13-week treatment in JNPL3 tauopathy mice (10 mg/kg intraperitoneally) robustly cleared insoluble tau (59-74% reduction), whereas soluble tau was not significantly affected. Considering that some form of equilibrium exists between these two pools of tau, the lack of effect on soluble tau may simply reflect continuous antibody-mediated transition of insoluble tau to soluble tau. A second in vivo confirmation of therapeutic benefits comes from the acute microdialysis study, in which a single injection of 5G2 (50 µg) reduced tau levels in brain interstitial fluid by 46% in the same JNPL3 model. In previous studies, using an acute treatment paradigm (2-3 injections of 4E6 tau antibody in htau and JNPL3 tauopathy models, soluble tau was primarily being cleared (Congdon et al., 2016;Wu et al., 2018;Wu et al., 2020). Although these studies cannot be directly compared because the antibodies are not the same, it seems logical that a chronic vs acute treatment would affect insoluble vs soluble pools of tau, respectively.

Neuronal Ca^2+^ dysfunction has for many years been implicated in AD (2017;Bezprozvanny and Mattson, 2008), and increased Ca^2+^ influx does have a close link to tau pathology (Furukawa et al., 2003;Mattson, 1990;McKee et al., 1990;Nixon, 2003). Neuronal calcium profile has been examined to some extent in tauopathy mouse models with most of the studies reporting functional abnormalities (Busche et al., 2019;Decker et al., 2015;Kopeikina et al., 2013;Kuchibhotla et al., 2014;Marinkovic et al., 2019;Overk et al., 2015;Wu et al., 2020). We have previously shown that tauopathy-induced Ca^2+^ dysfunction is most prominent in awake active mice and that imaging these animals before and after acute treatment can reveal its beneficial effect on neuronal function. The positive effects of the 5G2 tau antibody in the PS19 x Thy-1^GCaMP6^ mice were comparable to our findings with the 4E6 tau antibody against the P-Ser396, 404 region of tau in the JNPL3-AAV-GaMP6s model, in particular under running conditions (Wu et al., 2020). In both of these studies, analyzing the neuronal calcium profile under running conditions is more sensitive than during resting state to detect differences between wild-type and tauopathy mice, and for appreciating functional benefits of tau antibody treatment.

Overall, this study indicates robust benefits of targeting pathological tau truncated at Asp421 to prevent tau neurotoxicity, and to promote tau clearance in three different culture models as well as in two mouse tauopathy models under chronic and acute conditions. These positive effects are associated with prevention of tau-induced microgliosis in culture, reversal of microgliosis in vivo and functional in vivo benefits based on neuronal calcium imaging. Mechanistically, acute antibody-mediated tau clearance appears to be linked to the high affinity intracellular Fc receptor, TRIM21, which presumably promotes proteasomal clearance of the antibody-tau complex. On the other hand, long-term tau clearance is likely to be mainly caused by the antibodies’ ability to disassemble tau aggregates within the lysosomes and thereby facilitate their enzymatic degradation.

## Acknowledgements

This work was supported by the National Institutes of Health (NIH; NS077239, AG032611 and AG069475 for EMS), and Alzheimer’s Association (AARFD-22-926379 for AMA). We thank Dr. Peter Davies (Albert Einstein College of Medicine and Long Island Jewish Medical Center, Litwin-Zucker Research Center at Feinstein Institutes for Medical Research) for the tau antibodies PHF1 and CP27, which were subsequently obtained from Drs. Philippe Marambaud and Jeremy Koppel at Feinstein. We acknowledge the use of tissues procured by the National Disease Research Interchange (NDRI) with support from NIH grant 2 U42 OD011158.

## Ethics approval

Any data derived from human tissue did not include any patient identifiers. This study was conducted in accordance with US ethical guidelines and was deemed exempt from ethics approval by the Institutional Review Board (IRB). All mouse experiments were performed under an institutional animal care and use committee (IACUC) approved protocol with the mice housed in Association for Assessment and Accreditation of Laboratory Animal Care (AAALAC) approved facilities with access to food and water ad libitum.

## Availability of data and materials

All data needed to evaluate the conclusions in the paper are present in the paper and/or the Supplemental Materials.

## Authors’ contributions

AMA performed all aspects of the two-photon studies, including maintenance and crossing the colonies, surgical preparations, imaging and related analyses, as well as brain analyses of those brains by western blots and immunohistochemistry, with assistance from MW, SRM performed the culture studies and related analyses with help from DBS for the TRIM21 knock-down experiment, HBR maintained some of the animal colonies and performed the chronic treatment study in JNPL3 mice and some of the related analyses with help from EEC who performed the western blots and related analyses. AD and LAS-B performed the microdialysis studies and related analyses. SK performed the binding studies, and YL maintained some of the animal colonies. EMS oversaw the design of the experiments and wrote the first draft of the article. All authors had the opportunity to edit the article. EMS supervised the project.

---

**Supplementary Figure 1.**
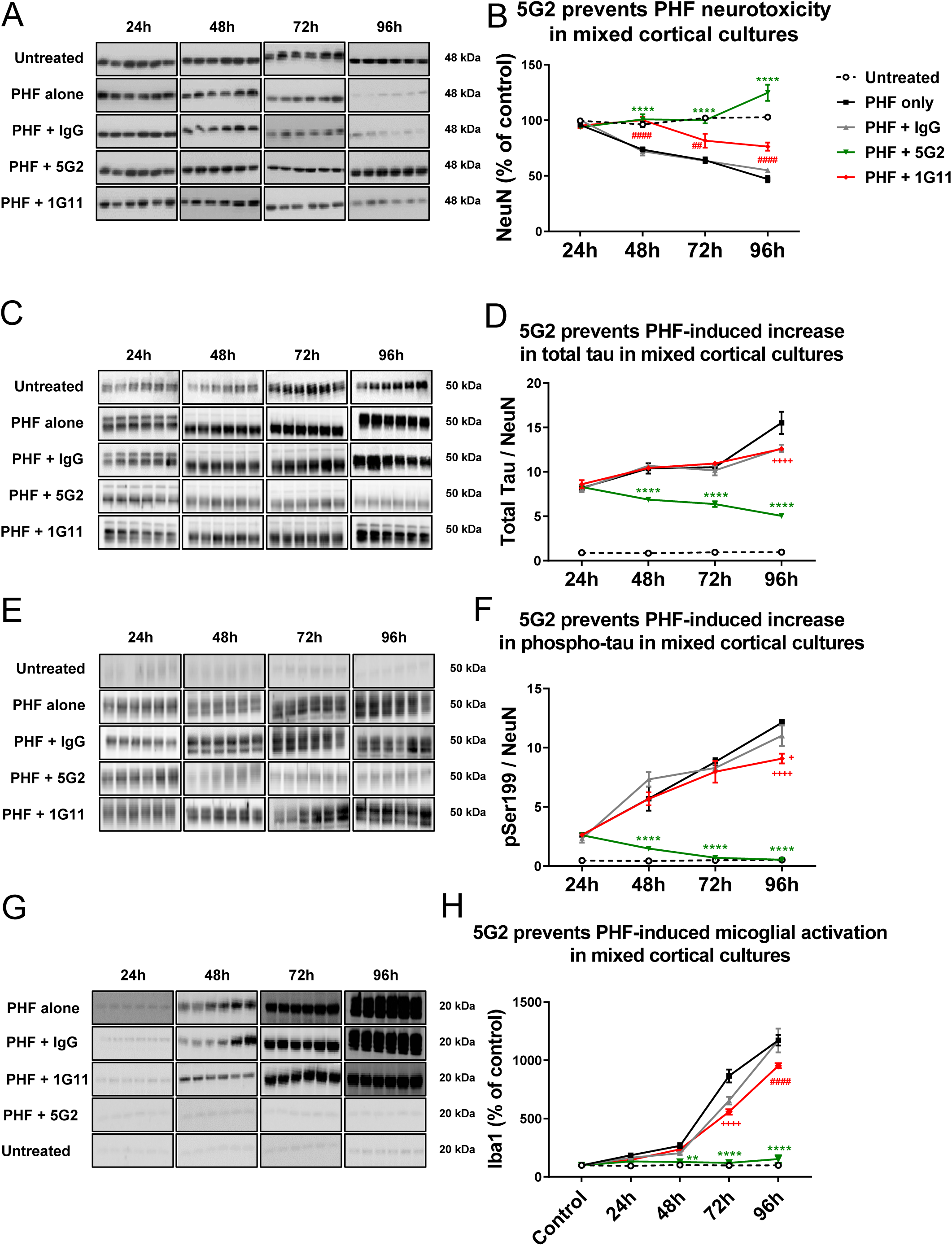
5G2 treatment prevented PHF-tau induced neurotoxicity, cleared total tau and phospho-tau, and prevented PHF-tau induced microgliosis in mixed cortical culture. Complete cortical culture from Day 0 P301L mice was co-treated with PHF (10 µg/ml) and Asp421 antibodies (5G2 or 1G11; 10 µg/ml) for a duration of 24 h, 48 h, 72 h and 96 h. Western blots were performed against NeuN, total tau, pSer199, and microglia (Iba1). Total tau and phospho tau levels were normalized against NeuN. **A, B)** A significant reduction in NeuN levels was observed in cells treated with PHF alone at 48 h (24%), 72 h (37%) and 96 h (54%), when compared to untreated cells, p < 0.0001. The control IgG co-treatment was ineffective in reducing this toxicity. 5G2 proved effective in completely blocking the neurotoxicity, wherein no significant alteration in the NeuN levels were observed from 24-72 h with an increase at 96 h (21%, p < 0.0001), compared to untreated cells. A partial prevention of PHF-induced neurotoxicity was observed with 1G11 co-treatment at 72 h and 96 h (20% and 26% reduction in NeuN, respectively), compared to the untreated cells (p < 0.001-0.0001), with complete prevention observed through 48 h. **C, D)** Total tau was increased several fold in the PHF-treated mixed culture. The 5G2 antibody gradually reduced it (48 h: 34%, 72 h: 39%, and 96 h: 68%, compared to cells treated with PHF alone, p < 0.0001 for all time points). The 1G11 antibody and control IgG were ineffective in reducing total tau levels at all time points except at 96 h where they reduced tau levels compared to PHF alone group (18-19% reduction, p<0.0001). **E, F)** Likewise, phospho-tau (p-Ser199) was increased several fold in PHF treated mixed culture. The 5G2 antibody gradually reduced it at 48 h (74%), 72 h (92%) and 96 h (96%), compared to cells treated with PHF alone, p < 0.0001 for all. These phospho-tau levels in the 5G2 treated group were not significantly different from those in untreated cells. 1G11 was slightly effective in reducing pSer199 tau at 96 h (p<0.0001 compared to PHF alone (25% decrease). **G, H)** Mixed cortical cultures treated with PHF alone, PHF + IgG and PHF + 1G11 had a robust increase in Iba1 levels at 24-96 h post-treatment compared to untreated mixed culture (PHF alone: 98%- 1074%, PHF + IgG: 74%%-1072%, PHF + 1G11: 54%%-855%, p < 0.01-0.0001). This PHF-induced microgliosis was modestly reduced by control IgG at 72 h (25%, p < 0.0001) and by 1G11 at 72 h (36%, p < 0.0001) and 96h (19%, p < 0.0001), compared to cells treated with PHF alone. However, the PHF-induced microgliosis was completely blocked by 5G2 co-treatment, resulting in comparable Iba1 levels to untreated controls. Statistical significance was determined by two-way ANOVA, followed by Bonferroni post hoc test for multiple comparisons, n = 6, values are represented as mean + or ± SEM, wherein **** p < 0.0001 PHF + 5G2 vs. PHF alone or PHF + IgG; ** p < 0.01 PHF + 5G2 vs. PHF alone; ####, ## p < 0.0001, p < 0.01 PHF + 1G11 vs. PHF alone or PHF + IgG; ++++ p < 0.0001 PHF + 1G11 vs. PHF alone, + p<0.05 PHF + 1G11 vs. PHF + IgG.

**Supplementary Figure 2.**
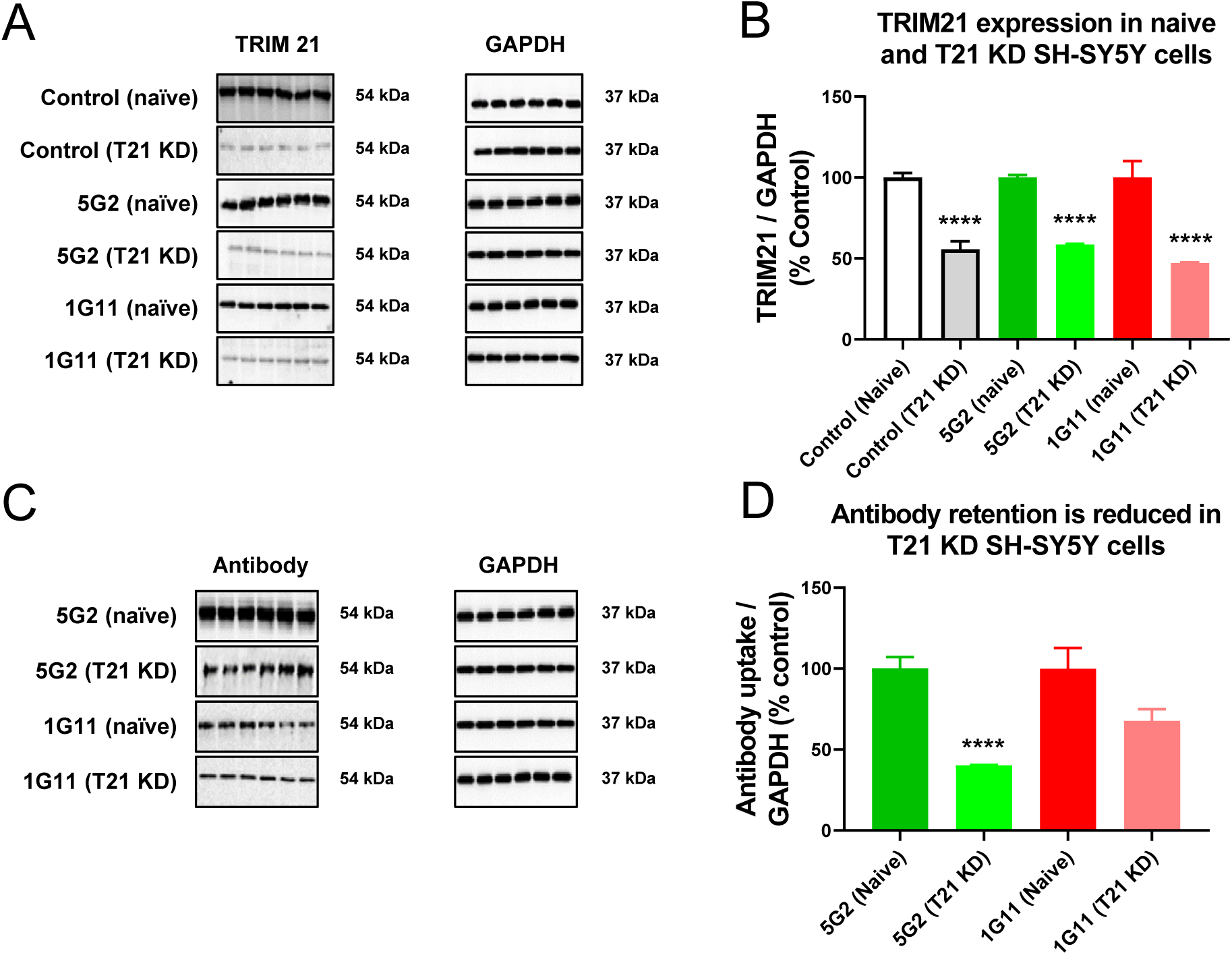
TRIM21 expression was associated with intracellular antibody retention. **A, B)** TRIM21 knockdown (T21 KD) resulted in 44% decrease in TRIM21 in differentiated SH-SY-5Y cells, compared to its naïve and otherwise identical control (p < 0.0001). Similar degree of knockdown (5G2: 41%, p < 0.0001; 1G11: 53%, p < 0.0001) was seen in the 5G2 and 1G11 antibody treated cells (5 µg/ml for 24 h), when compared to their respective naïve antibody treated cells. **C, D)** T21 KD reduced antibody retention. The antibody uptake was analyzed by probing Western blots with anti-mouse IgG1 HRP conjugate secondary antibody. A reduction in the uptake was observed for both the antibodies in the T21 KD cells (5G2: 60%, p < 0.0001 and 1G11: 32%, p = 0.069) when compared to the respective naïve antibody treated cells (shown as % of those controls). Statistical significance was determined by two-way ANOVA, followed by Bonferroni post hoc test for multiple comparisons, n = 6, values are represented as mean + SEM and normalized to GAPDH, which was comparable in the different groups (see Suppl. Figure 1).

**Supplementary Figure 3.**
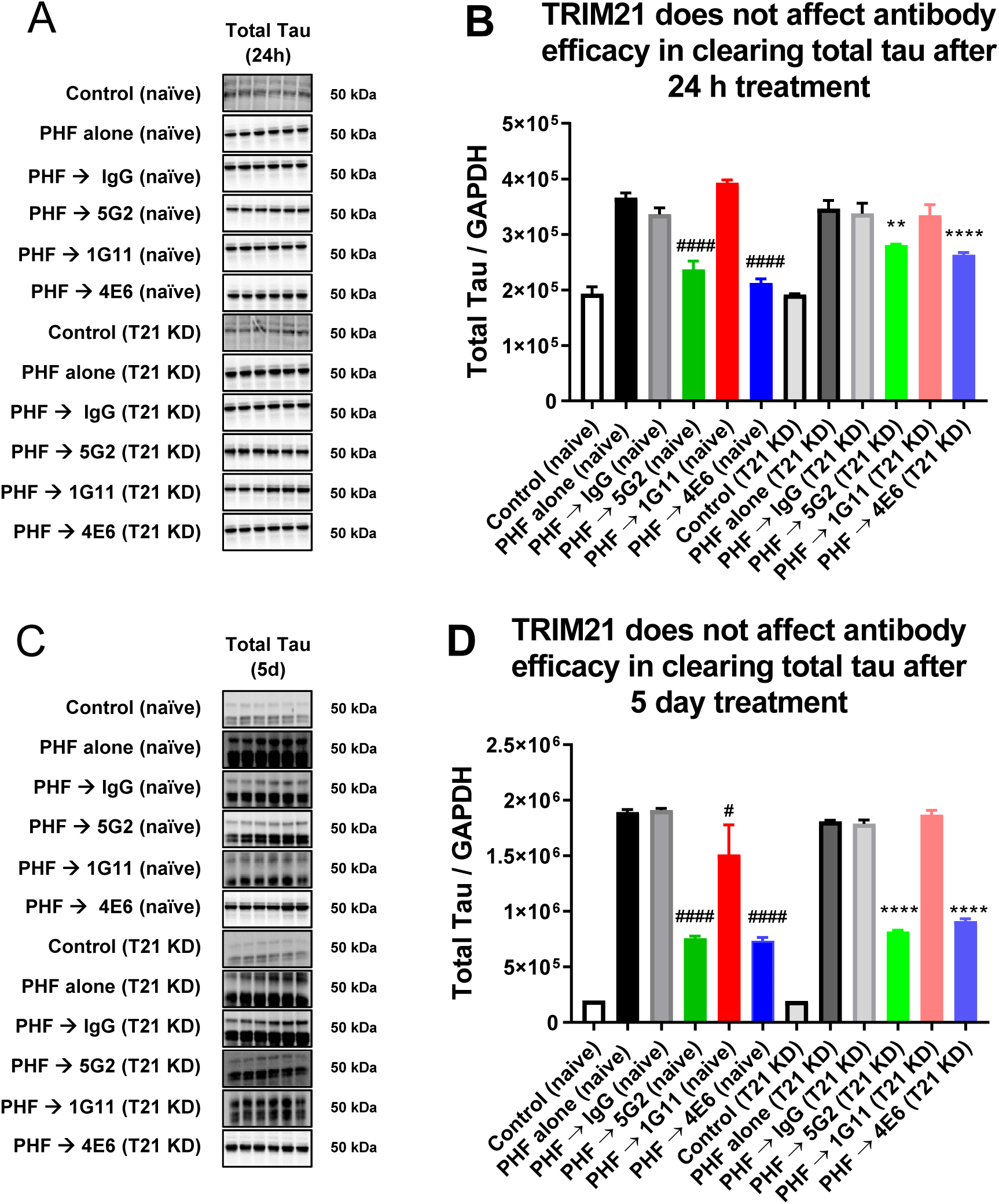
TRIM21 knockdown (T21 KD) influences antibody-mediated tau clearance to a varying degree, depending on antibody efficacy. Differentiated naïve and T21 KD SH-SY5Y cells were pre-treated with PHF (5 µg/ml) for 24 h, followed by treatment with antibodies, 5G2, 1G11 or 4E6 (5 µg/ml) for 24 h and 5 days, followed by western blots for total tau. **A, B)** At 24 h, 5G2 and 4E6 reduced total tau in naïve cells (5G2: 36%, p < 0.0001; 4E6: 42%, p < 0.0001) and T21 KD cells (5G2: 19%, p < 0.01, 4E6: 24%, p < 0.0001), compared to their respective cells treated with PHF alone. **C, D)** At 5 days, 5G2, 1G11 and 4E6 reduced total tau in naïve cells (5G2: 60%, p < 0.001; 1G11: 20% decrease, p < 0.05; 4E6: 61%, p < 0.0001) and 5G2 and 4E6 in T21 KD cells (5G2: 55%, p < 0.0001; 4E6: 50%, p < 0.0001), in comparison to their respective cells treated with PHF alone. Antibody-mediated reduction in total tau did not differ significantly overall between naïve and T21 KD at either time point, although post-hoc analyses revealed some subtle differences (24 h: 5G2, p = 0.056; 4E6, p < 0.05; 5 days 1G11, p < 0.05). in particular, the attenuation in tau clearance mirrored TRIM21 KD for the more efficacious antibodies, 5G2 and 4E6, under the acute 24 h treatment conditions as shown above (5G2: 36% to 19%; 4E6: 42% to 24%). Control IgG treatment was ineffective in both cell types at both time points. Statistical significance was determined by two-way ANOVA followed by Bonferroni post-hoc test for multiple comparisons, n = 6, values are represented as mean + SEM. #, ####: p < 0.05, p < 0.0001 in naïve cells compared to PHF alone. **, ****: p < 0.01, p < 0.0001 in T21 KD cells compared to PHF alone.

**Supplementary Figure 4.**
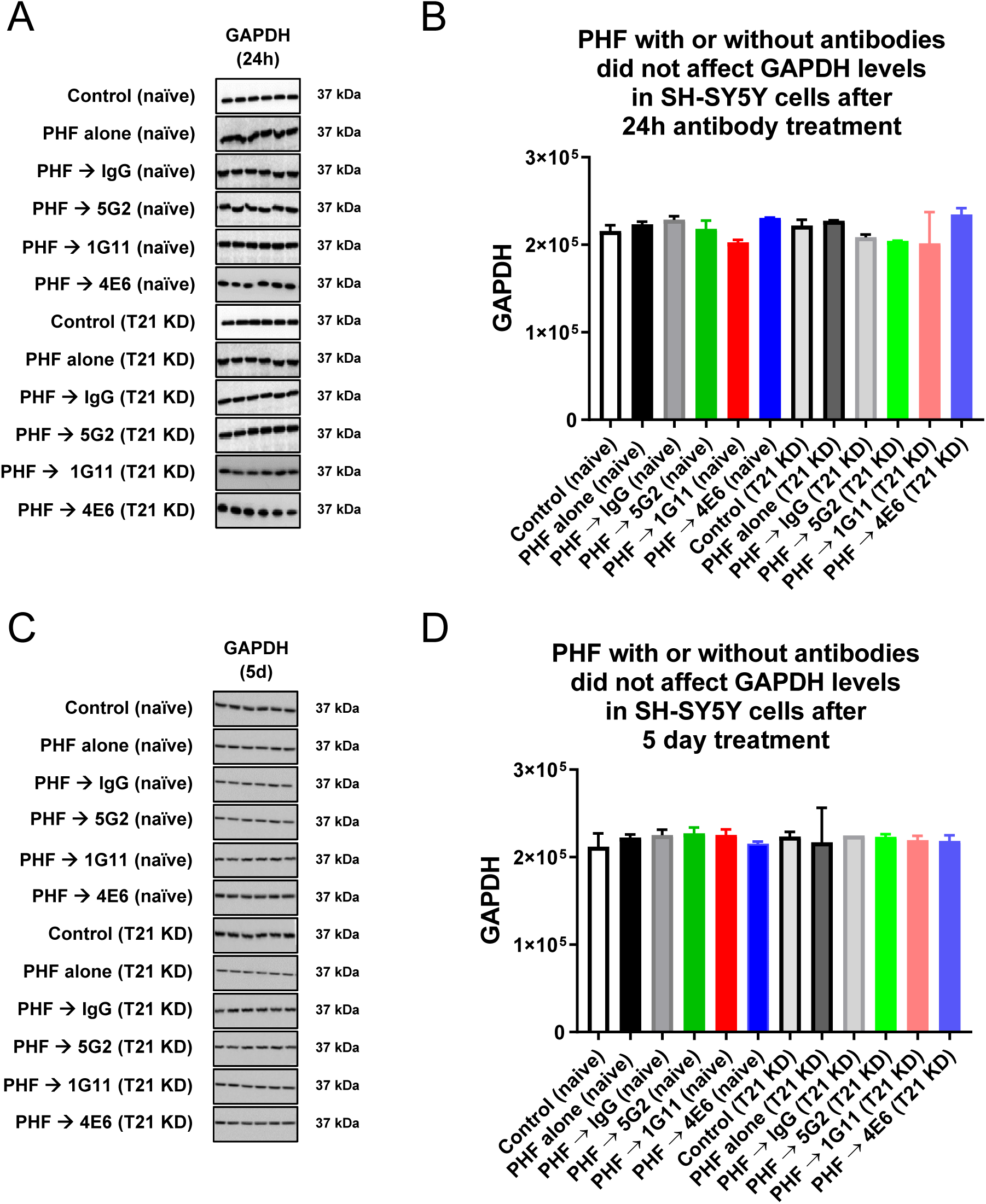
Treatment with PHF with or without tau antibodies did not affect GAPDH levels in naïve or TRIM21 knockdown (T21 KD) cells. Differentiated naïve and T21 KD SH-SY5Y cells were pre-treated with PHF (5 µg/ml) for 24 h, followed by treatment with antibodies, 5G2, 1G11 or 4E6 (5 µg/ml) for 24 h and 5 days, followed by western blots for GAPDH. These are the same cells that were analyzed for total tau in Figure 4. **A, B)** At 24 h, GAPDH levels were not changed in any of the groups. **C, D)** At 5 days, GAPDH levels were not changed in any of the groups. Statistical significance was determined by two-way ANOVA followed by Bonferroni post hoc test for multiple comparisons, n = 6, values are represented as mean + SEM.

**Supplementary Figure 5.**
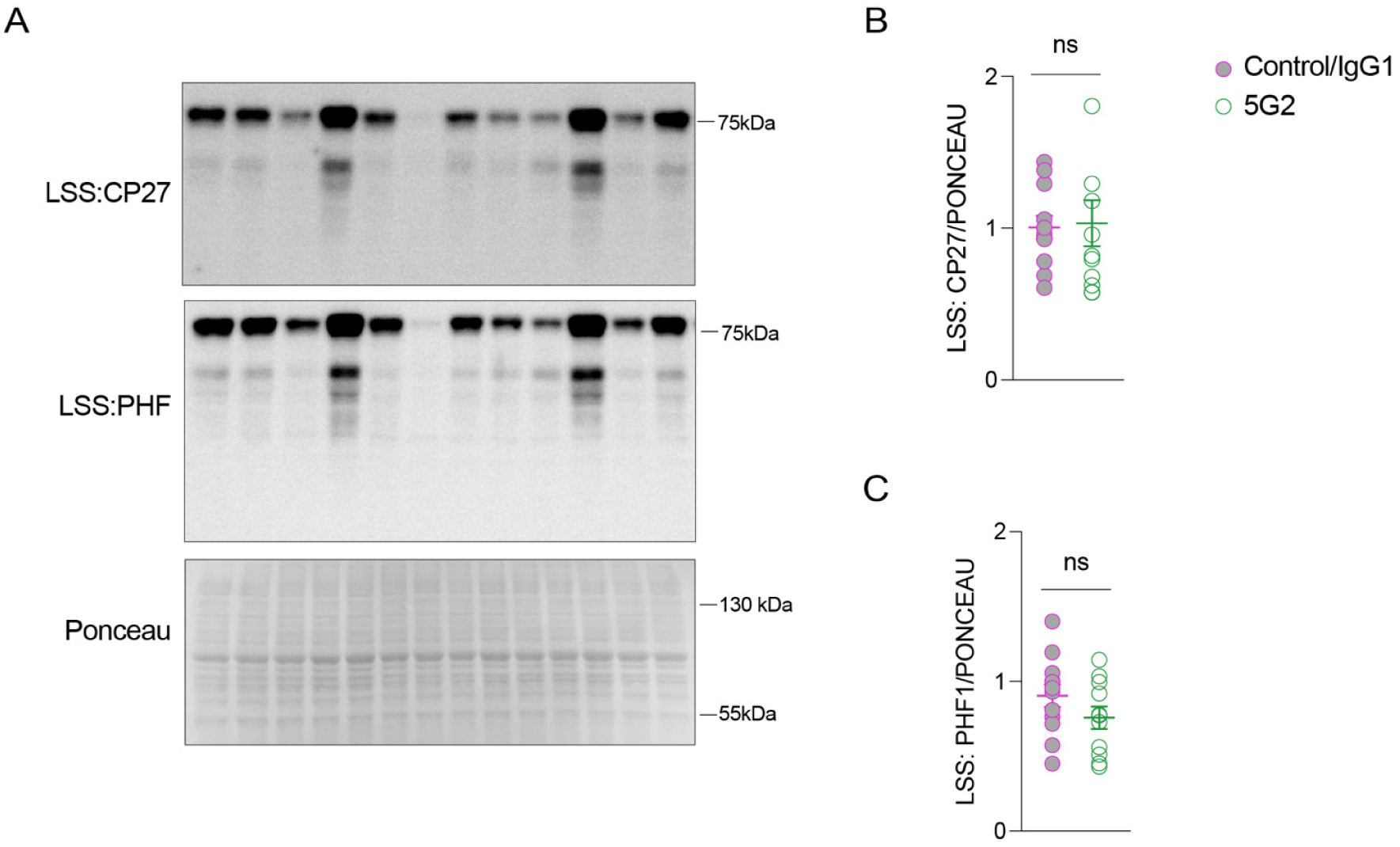
Acute 5G2 immunotherapy does not reduce CP27 and PHF-1 signal in soluble fraction of brain homogenate. **A)** A representative blot showing CP27 and PHF-1 signal in soluble brain fraction (n = 5 PS19 mice (control), n = 5 IgG-injected PS19 mice, n = 11 5G2-injected PS19 mice. **B, C)** Quantification of CP27 and PHF-1 signal.

